# Contrastive learning for cell division detection and tracking in live cell imaging data

**DOI:** 10.1101/2024.08.16.608296

**Authors:** Daniel Zyss, Amritansh Sharma, Susana A Ribeiro, Claire E Repellin, Oliver Lai, Mary J C Ludlam, Thomas Walter, Amin Fehri

## Abstract

Fluorescent live-cell microscopy is essential for understanding cellular dynamics by imaging specific molecules, their interactions, and biochemical states in live samples. It is crucial for biological research and drug screening. However, live-cell imaging often requires balancing temporal resolution with cell viability (i.e. conditions enabling cells to thrive) because of photo-toxicity. Consequently, there is increasing interest in lowering temporal resolution to allow for extended observation periods and to study event sequences that uncover causal relationships and mechanistic insights. Tracking cells in video microscopy with low temporal resolution remains a challenge. We introduce a new integrated methodology that uses contrastive learning and graph-based approaches to improve cell division detection and tracking. Our approach employs contrastive learning models to create cell representations that facilitate the detection of cell divisions and enhance cell tracking. Of note, we propose a weakly-supervised constrastive learning approach to build robust temporal cell representations through time-based augmentations. Additionally, we introduce an innovative graph optimization technique to identify cell tracks based on these representations and observed division events. We evaluate our methods on an in-house dataset and public datasets from the Cell Tracking Challenge, achieving substantial performance improvements in both native and reduced temporal resolutions. Our methodology thus enhances adaptability to various temporal resolutions, improving precision and efficiency in live-cell microscopy analysis. This advancement is particularly beneficial for extended drug screening studies, ensuring cell viability and maintaining normal cell homeostasis, which is vital for therapeutic research.

## 1. Introduction

The study of individual cells’ phenotypes over time is pivotal in understanding the cellular dynamics and responses to drug interventions, particularly in disease-relevant model systems. This time-resolved single cell phenotyping indeed offers critical insights into the mechanisms of action (MoA) of drugs, enabling a deeper comprehension of cellular and signaling events in disease responses to treatment. Furthermore, the integration of single-cell behavioral data from live-cell imaging with genomic sequencing is anticipated to revolutionize the identification of novel biomarkers and actionable targets, especially in the realm of cancer drug discovery by helping better identify compounds’ MoA as well as model and forecast resistance to treatment [1].

In drug screening live-cell studies, achieving a balance between detailed readouts and minimal perturbation to the model system is crucial. Prolonged exposure to fluorescent light can induce phototoxic effects, impacting cell viability and causing non-physiological behaviors. Therefore, a choice arises: either limit the experiment duration, sacrificing long-term phenotypic insights, or reduce the temporal resolution. Choosing the latter results necessarily in time-lapse experiments with low temporal resolution, which pose challenges in processing due to significant morphological and phenotypic changes between frames [2]. This gap complicates automated cell tracking, as traditional tracking methods (usually based on spatial distance and overlap) struggle to accurately follow cells through large time intervals. Figure 1(A) shows the various trade-offs encountered through the experimental set-up and the image acquisition constraints. The impact on division detection and tracking of running longer experiments with lower temporal resolution is illustrated in Figure 1 (B).

**Figure 1.**
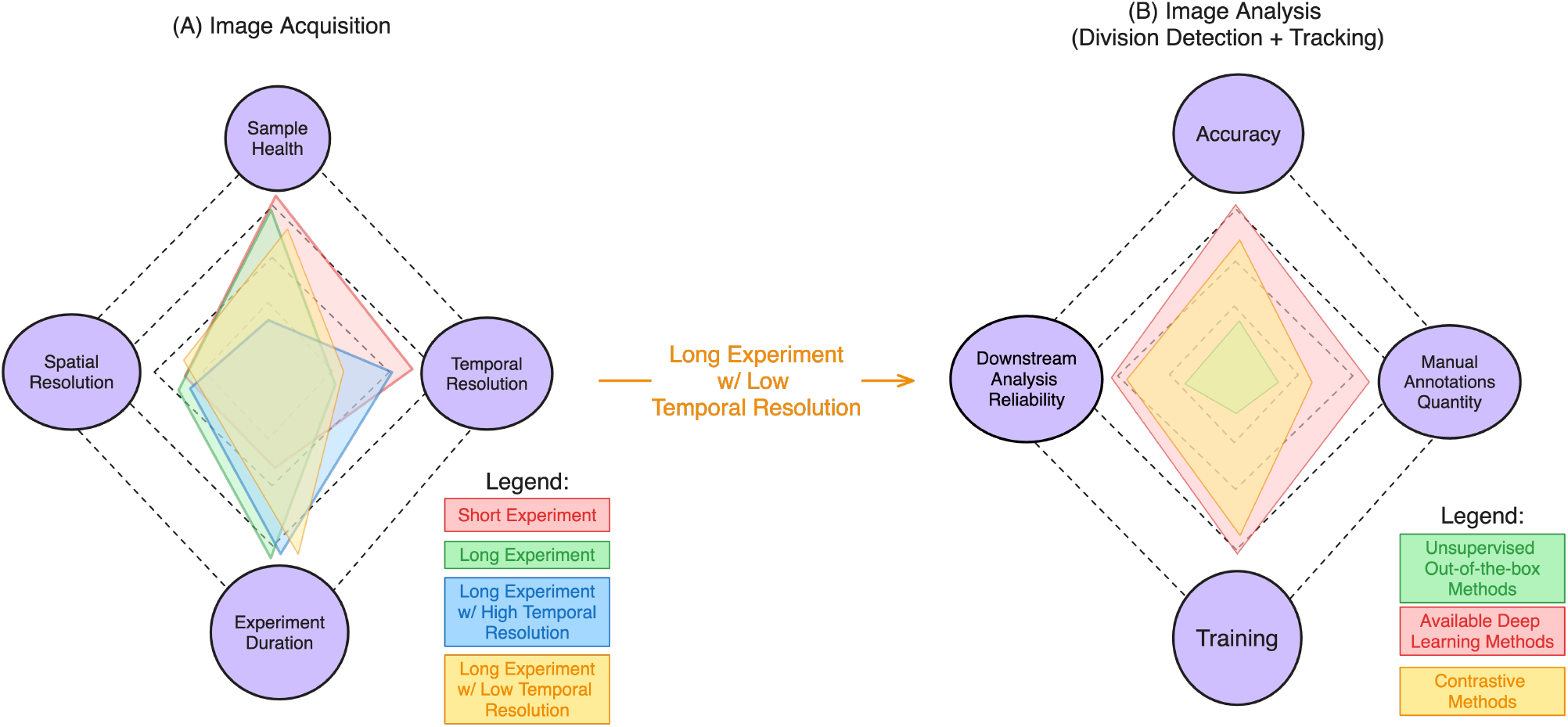
Acquisition and analysis trade-offs in video-microscopy: **A)** This diagram explores the compromise we must navigate between the length of the experiment and the temporal resolution during live cell imaging. To observe cells for longer periods, we must reduce the temporal resolution to protect sample health, largely due to the risk of photo-toxicity. On the flip side, to achieve high temporal and spatial resolutions, we must limit the duration of the experiment to ensure the quality of live fluorescent imaging. **(B)** In the context of studies involving live cells over extended periods with low temporal resolution, the strategies for detecting cell divisions and tracking involve significant trade-offs to secure reliable downstream analysis. These strategies range from those demanding extensive manual annotations and training for higher accuracy to simpler, less demanding methods with reduced accuracy. In this work, we present a novel method that employs contrastive learning to reduce the need for manual annotations while still achieving high accuracy levels, thereby ensuring the reliability of cell tracks downstream analysis. These spider-plots are inspired by [3].

Cell tracking techniques generally fall into two categories [4]: supervised deep learning methods [5, 6] and classical assignment methods based on hand-crafted features [7]. Both approaches are highly sensitive to variations in arbitrarily selected hyper-parameters, experimental conditions, and imaging setups. As temporal resolution decreases, spatial proximity and movement become less powerful at establishing the correct temporal correspondences. To close the gap, one can turn to using morphological resemblance between instances of the same cell in consecutive time-frames, a task deep learning usually excels at. Indeed, such learning-based methods have the advantage to directly learn from the data potentially more complex rules for finding associations. However they demand extensive annotation and training efforts that must be repeated every time experimental conditions vary. Our research addresses these limitations and enables the swift adaptation of computational analysis pipelines to such changes, particularly in the context of low temporal resolution imaging.

In this manuscript, we introduce and validate an innovative framework for cell division detection and cell tracking in fluorescence video microscopy, able to handle low temporal resolution datasets, while requiring limited manual annotation. The division detection utilizes a semi-supervised approach incorporating elements of self-supervised learning [8], while the tracking algorithm combines dynamic programming with cell similarity estimations derived from both hand-crafted and annotation-free weakly-supervised contrastive deep learning features. This dual approach accommodates the important morphological and phenotypic changes characteristic of low temporal resolution microscopy. By addressing the trade-off between experiment duration and temporal resolution, our method provides a flexible and robust tool for comprehensive cell analysis in long-term studies, opening new avenues for understanding complex biological processes.

We evaluate our methods on an in-house dataset, as well as benchmark them on datasets sourced from the Cell Tracking Challenge (CTC) [9, 4]. Additionally, we provide access to in-house annotated datasets to the wider research community. Our results showcase that our framework achieves state-of-the-art performance in both cell division detection and tracking across diverse cell lines and under varying experimental conditions. Remarkably, it accomplishes this with minimal annotation requirements. Consequently, our framework emerges as a valuable asset for live-cell drug screening and more generally live-cell imaging and cell biology research.

## 2. Related work

### 2.1. Cell division detection

The field of cell division detection in microscopy has predominantly concentrated on identifying mitotic phases in order to monitor mitotic progression i.e. the process of cell division in which a single cell divides to produce two genetically identical daughter cells. These methods usually operate on datasets with fluorescent markers reporting on chromosome segregation or the mitotic spindle, or structural markers which can be leveraged to detect the different stages of division. They are generally based on a sequential workflow, consisting in segmentation, feature extraction and classification [10–14]. Other methods such as [15, 16] utilize dynamic changes observed in video frames to identify mitotic patterns. Techniques introduced by [17], [18], and [19] employ hidden conditional random fields to identify mitotic event patches in images, but these methods demand high temporal resolution as well. Developed specifically for phase contrast microscopy, they might furthermore not provide robust models for the phenotypic dynamics encountered in fluorescent microscopy, even at high temporal resolutions. All these approaches require a high time resolution which is sufficient for at least one mitotic figure to be observed, e.g. MitoCheck [10] uses a time resolution of 30 min. Recent developments have leveraged deep learning techniques for division detection. Mao et al. [20] introduced a Hierarchical Convolutional Neural Network (HCNN) specifically for time-lapse phase contrast microscopy, achieving high accuracy with a annotated training dataset of 1,436 images at relatively high temporal resolution. Building on this, Mao et al. [21] enhanced their methodology by integrating Low Rank Matrix Recovery to isolate division candidates, subsequently classifying and analyzing them with an HCNN followed by a Long-Short Term Memory (LSTM) model, showing excellent detection accuracy on phase microscopy images at a 5-minute temporal resolution. This trend of specialized division detection methods is further discussed by Liu et al. [22]. A notable innovation is the combination of division detection within tracking algorithms, exemplified by the work detailed in [23], which employs deep learning for candidate selection in tracking and division detection, along with graph optimization using hand-crafted features for tracking. This approach effectively but not entirely addresses the challenges posed by low temporal resolution in fluorescent microscopy datasets from the Cell Tracking Challenge (CTC), managing to work with resolutions up to 30 minutes as discussed by Maka et al. [24].

Most division detection and tracking methods have thus been developed for such high temporal resolution, supported by standard tracking benchmarking datasets like the Cell Tracking Challenge (CTC) dataset [4] which temporal resolutions range from 5 to 30 min. At lower temporal resolutions the risk increases of missing mitotic figures and identifiable dynamic patterns that are crucial for identifying cell division events with these methods. Moreover, merely recognizing an imminent cell division does not address the challenge of accurately linking daughter cells to their respective mother cells. In this paper, we address the challenge of *semantic division detection*, i.e. the process of recognizing division events where one mother cell in a frame transitions to its two daughter cells in the next frame, regardless of division associated phenotypes and related fluorescent reporters. Lineage Mapper [7] and Ilastik [25] are commonly employed methods for semantic division detection using segmentation masks. Both methods uses logic-based algorithm to deduce division from segmentation maps using simple criteria like roundness, overlap or spatial distance. They however struggle with low temporal resolution or experiments involving highly dynamic morphological changes, leading to inadequate detection.

The existing methods for division detection and tracking do not meet the demands of live-cell drug discovery over prolonged experimental duration. There is thus a need for new approaches able to interpret a broad spectrum of morphological and phenotypic changes associated with cell division and that remain effective even when temporal resolution decreases beyond 30 minutes. Furthermore, deep learning-based methods often require a high volume of annotated data, which limits their adoption in practical settings.

### 2.2. Cell tracking

Cell tracking is an important and active field of bioimage analysis. A plethora of different methods have been applied to track individual cells over time. Given the challenge of categorizing these into a clear taxonomy and the extensive literature already available, it is impractical to enumerate all developments in cell tracking algorithms here. Instead, we direct readers to Ulman et al. [9], which provides a comprehensive comparison and evaluation of various cell tracking algorithms using the CTC dataset. Noteworthy among the methods reviewed are: (1) a linking by detection approach by Turetken et al. [26], which combines Integer Programming with Ellipse fitting, and (2) a model-based tracking strategy by Magnusson et al. [27], employing the Viterbi algorithm to iteratively optimize a probabilistic segmentation-based motion model.

As many fields in bioimage analysis, cell tracking has been impacted by the advent of deep learning. [28] provides a comprehensive review of the advancements in machine-learning enhanced cell tracking, highlighting the increasing role of deep learning in the field. Accordingly, in the last decade, several cell tracking challenges have been organized, aiming to benchmark the performance of different tracking algorithms under various conditions [4]. These challenges have greatly contributed to our understanding of the strengths and weaknesses of current approaches.

One key insight is the difficulty of dealing with low temporal resolution datasets, where significant cell morphological changes and cell displacement can occur between frames [6]. Indeed, many of these advanced methods such as ALFI [29] and CycleBase [30] require high temporal resolution data to be effective, which is not always feasible due to the limitations imposed by phototoxic effects in live-cell imaging, as illustrated in Figure 1 and discussed in Section 1. Similarly, the weakly-supervised learning and graph-based approach proposed by Hirsch et al. [31] for tracking whole-embryo C. elegans lineages may also have limited applicability to datasets featuring important morphological changes and low temporal resolution.

Furthermore, these state-of-the-art [4, 29, 30] methods rely heavily on manual tracking annotations, which represent a significant burden and constraint in general and in particular in the context of drug screening due to the associated versatility of changes in experimental variables (cell-lines, fluorescent reporters, imaging conditions, …) often within the same study.

### 2.3. Self-supervised and weakly-supervised contrastive learning

Contrastive learning is a powerful metric-learning paradigm. It relies on the principle of contrasting samples against each other to learn attributes that efficiently discriminate data classes. Self-supervised contrastive learning works by learning by self-comparison, i.e. by encouraging the model to identify similarities and differences across various augmentations of the same image. This approach has shown remarkable efficacy in learning rich, robust representations from unlabeled data, a notable example being the simCLR framework [8].

Weakly-supervised learning is an umbrella term covering studies aimed at learning with incomplete, inexact or inaccurate supervision, possibly obtained in an automated fashion. Zhou’s review [32] provides an overview of weakly-supervised learning, underscoring its relevance in scenarios with limited expert-annotated data.

The combination of weakly-supervised and contrastive learning paradigms has emerged as a potent strategy for deriving meaningful representations from data, particularly when faced with the challenge of limited expert annotations. This approach diverges from traditional self-supervised contrastive learning, where augmentations of the same image are utilized, by embracing weak supervision techniques. Theoretical advancements, such as those proposed by Cui et al. [33], underscore the efficacy of integrating semi-supervised learning and learning with noisy labels into the contrastive learning framework, demonstrating superior performance in downstream tasks compared to standard self-supervised methods. Similarly, research by Zheng et al. [34] refines this concept by adapting the simCLR architecture to accommodate weak classifier labels, thereby enhancing image representation robustness with minimal labeled data and mitigating class collision issues. Tsai et al. [35] further expand upon this theme, employing auxiliary data attributes to generate weakly-supervised relationships between data samples, thereby augmenting the performance of downstream models and surpassing traditional self-supervised methods. A particularly relevant application of these concepts is observed in the field of microscopy, as Borowa et al. [36] have innovatively applied weakly-supervised contrastive learning techniques to cell classification in drug screening studies. Their approach utilizes a linear model on basic morphological features and leverages image experimental metadata, such as compound concentration, for effective cell embedding and classification with minimal annotations. This methodology exemplifies the potential of weakly-supervised contrastive learning in facilitating efficient drug screening analysis processes.

In this paper, we harness the potential of both self-supervised and weakly-supervised contrastive learning to address the challenges of cell division detection and cell tracking, in a field often hampered by the scarcity of annotated data and the diversity inherent to biological datasets. Our approach aims to provide an efficient solutions applicable across various research contexts, while minimizing the dependency on extensive annotated data.

## 3. Materials and methods

### 3.1. Data

Our study leverages live-cell video microscopy data derived from High Content Screening (HCS) assays on the A375 cell line. Live-cell HCS assays consist in affecting a disease model (i.e. a cell line) with multiple perturbations (e.g. a drug, a mutation, etc.) and systematically quantifying their impact over time by analyzing the resulting video-microscopy data to help answer a biological question and/or assist drug discovery efforts. The A375 cell line, a human malignant melanoma line, is extensively used in cancer research due to its well-characterized nature and responsiveness to various treatments [37]. In our assays, this cell line was genetically modified to express several fluorescent reporters of the MAPK pathway [38]: mCerulean-RAF, Venus-RAS, mCherry-ERK, and miRFP670-MEK. These reporters allow for the detailed observation of key signaling pathways involved in cell proliferation and response to drug treatments. We plated 1000 cells/well in a 384 well plate, and treated them with 9 point half-log dilution of 6 compounds and DMSO control. The cell populations in our assays were subjected to diverse treatment regimens, including changes in drug types and concentrations, providing a comprehensive dataset to observe a range of cellular behaviors and responses.

We imaged cells at 4 hours intervals for 72 hours starting 10 minutes after initial treatment. We chose this extended intervals to minimize phototoxicity in order to study those cells over extended period of time and to adapt to the constraints of long-term imaging in HCS assays. This low temporal resolution presents unique challenges for cell tracking and division detection, as outlined in sections 1 and 2.

In the context of our research, we annotated some videos of our A375 dataset to train and evaluate our division detection method and to evaluate our cell tracking method. Specifically, the division dataset comprises 261 annotated division events across 10 videos. These annotations capture a diverse array of division scenarios, providing a comprehensive basis for training and testing our division detection algorithm. The tracking dataset encompasses 528 distinct cell tracks, for a total of 1538 individual cell observations across time within 3 videos for which we also have annotated division events. We designed that dataset to reflect the dynamic nature of cell movement and interaction over extended periods, which is critical for assessing the performance and accuracy of our unsupervised tracking method. For each of those image sequence we also relied on previously acquired segmentations. Our dataset was segmented using a segmentation workflow presented in a previous work [39]. We are publicly releasing this annotated dataset. It can be found in the following repository: https://figshare.com/articles/dataset/Division_Detection_and_Tracking_Annotated_Dataset/25766259.

To benchmark our method, we also use datasets from the Cell Tracking Challenge (CTC), a well-established resource providing a variety of cell tracking scenarios and datasets [24]. While the CTC datasets typically feature much shorter time intervals between frames, we adjusted these datasets to match the lower temporal resolution of our A375 cell line dataset by removing intermediate frames. This adjustment ensures a consistent basis for evaluating the effectiveness of our tracking and division detection methods under similar temporal resolution constraints, thus allowing a direct comparison and validation of our methodology against established benchmarks. We rely on the CTC ground truths for cell segmentation.

All datasets are showcased in Figure 2, showing the first 4 images for each sequence so as to reflect their different evolution speed.

**Figure 2.**
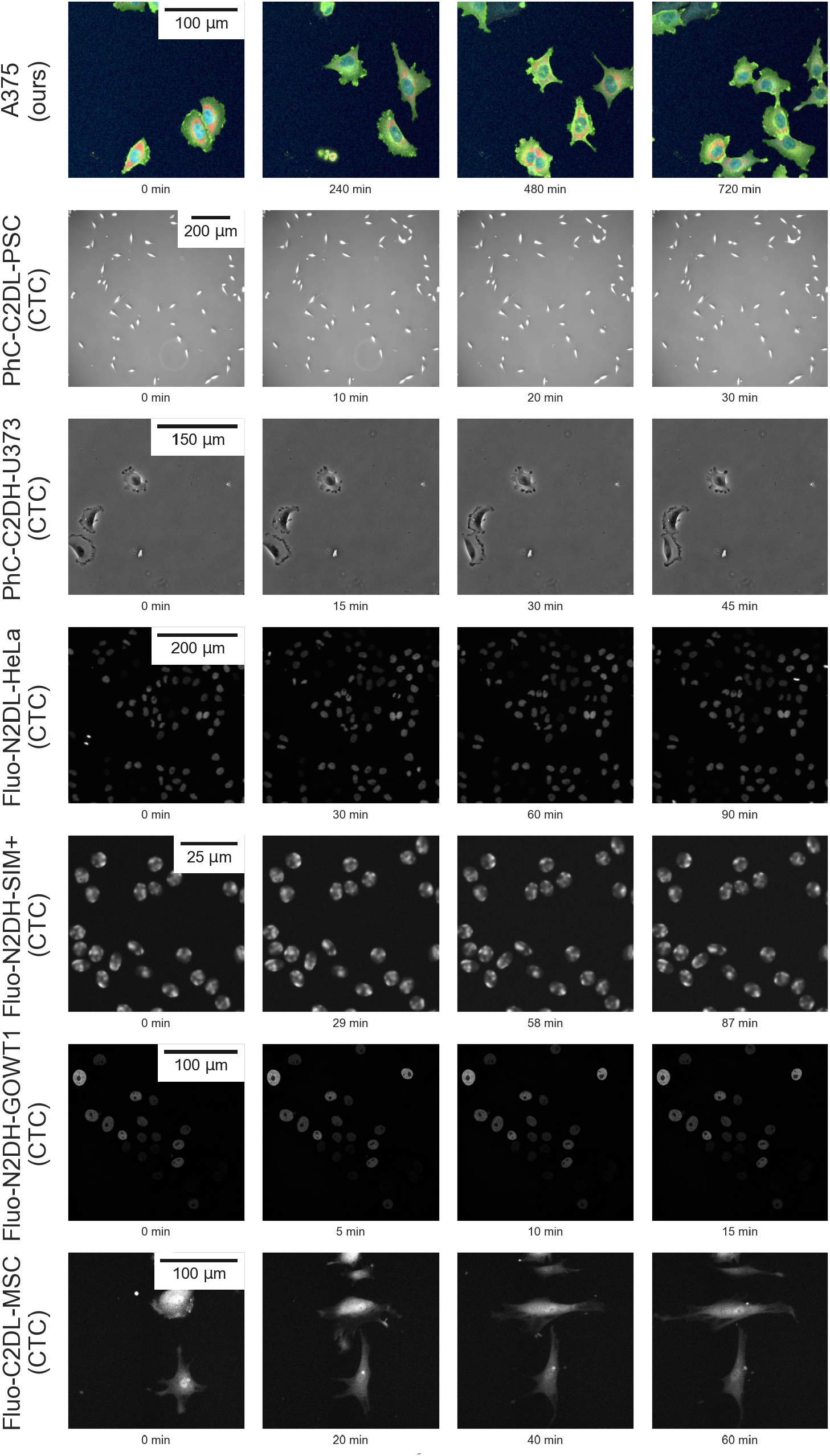
Time-lapse examples extracted from both our dataset and the CTC datasets. We display the initial four time-points of a single video for each cell line. Additionally, we provide the timestamps corresponding to each image capture along with the time scales of the respective datasets for comparison. The A375, PhC-C2DL-PSC and Fluo-N2DH-SIM+ images are zoomed in for clarity.

### 3.2. General workflow

The general workflow presented in this paper consists in two modules: (i) first we use a cell division detection module to extract cell division events, (ii) then we use a tracking module that takes as inputs these cell division events to derive cell tracking and cell lineage. Hereafter we describe these two modules, and their respective frameworks - in particular how they are trained and used at inference. Both frameworks are structured around an upstream pre-text task, aimed at deriving representations for individual cells, and a downstream target task, which utilizes these representations to facilitate division detection or tracking.

The division detection module identifies mother-daughter cell triplets in videos while requiring minimal human annotations. To do so, we develop a robust representation for these triplets, enabling a classifier to learn effectively from few annotations and distinguish positive division events in highly imbalanced datasets. This division detection training and inference workflow is illustrated in Figure 3. The training process comprises three stages: (1) Train a dedicated self-supervised learning model, designed to learn representations for cell image patches and their temporal context; (2) Identify mother-daughter candidate triplets in videos by selecting triplets of cells which conform to our criteria, using the annotated videos we label the candidates corresponding to division events as positive samples and the others as negatives for subsequent training; (3) Train a classification model to detect division events, using the candidate representations from the annotated videos. The inference for a previously unseen video then comprises three steps: (1) Build a graph structure of its cells, adhering to the same constraints employed during the training phase; (2) Generate representations for every cell within this graph, and then classify each triplet’s representations using the classifier to distinguish division events and accordingly label the graph; (3) Extract division events from the graph to ensure the coherence and uniqueness of each division event and mother-daughters relationship.

**Figure 3.**
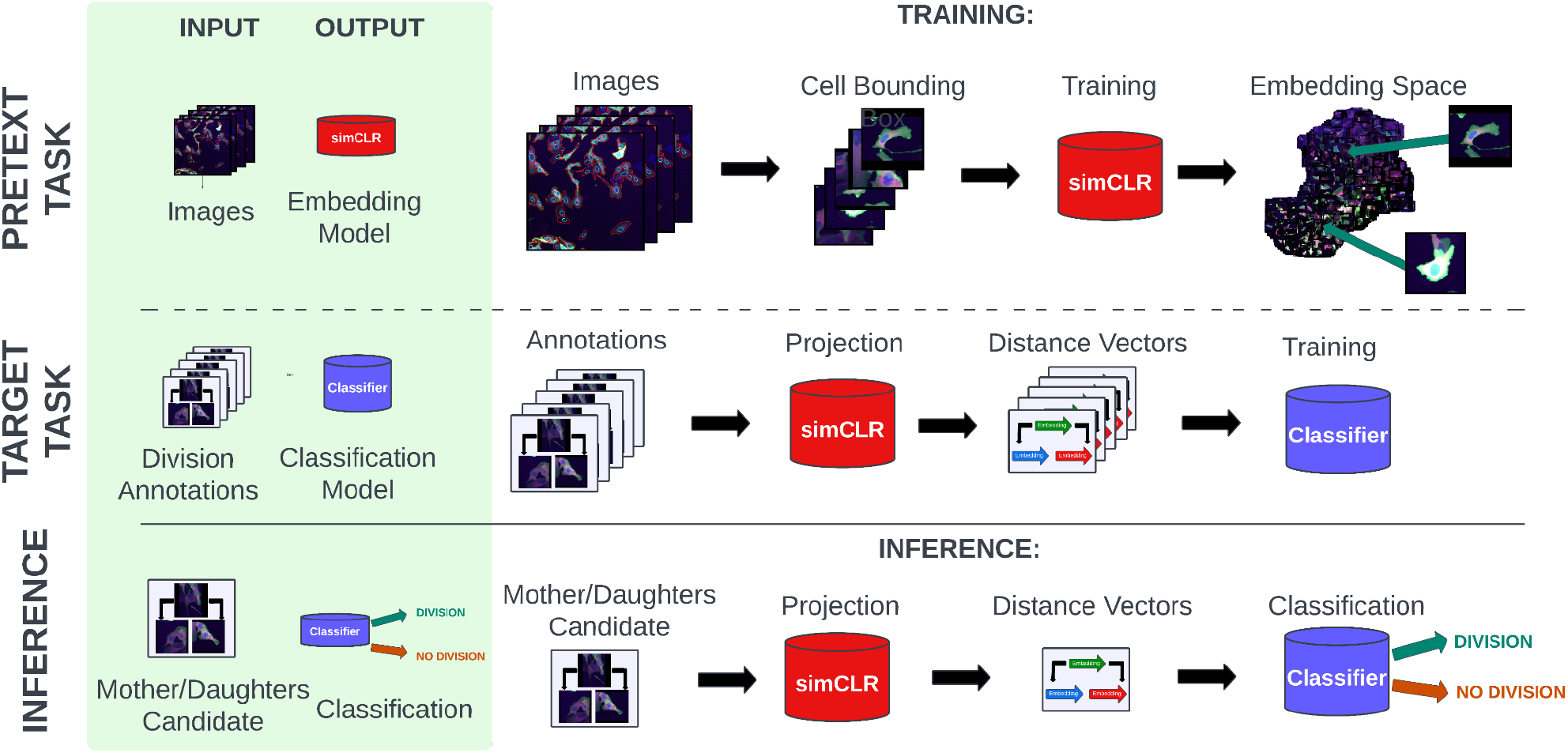
Division detection workflow: Our method’s training phase (top “training” section) comprises two stages: (1) The pretext task involves training an embedding model (here, a simCLR model) with patches extracted from individual cells’ Regions of Interest; (2) The target task involves training a classification model that utilizes features from the embedding model to detect division events. The inference phase on a new video (bottom “inference” section) involves three steps: (1) Selection of mother/daughter candidates from the video based on predefined constraints; (2) Generation of learned (simCLR) representations for each candidate triplet using our trained model; (3) Classification of each candidate into negative or positive division events using the trained classifier.

The cell tracking module aims at following individual cells over time, even at low temporal resolution, where spatial proximity is not the only relevant feature to consider. To do so, it uses weakly-supervised contrastive representation learning to generate robust features and graph assignment optimization. The tracking training and inference workflow is illustrated in Figure 4. The weakly-supervised contrastive model training comprises two main steps: (1) Generate “weak” preliminary tracks with an unsupervised algorithm, providing useful cell match pairs for further training despite their imperfections; (2) Leverage these weak examples to train a contrastive model and learn cell representations over time. Inference is then a two-step process: (1) Use the weakly-supervised contrastive model to extract features for each cell, enabling precise cell identification and tracking across frames; (2) Use graph optimization techniques that integrate spatial and other relevant features with our tracking contrastive features to construct accurate cell sequences over time.

**Figure 4.**
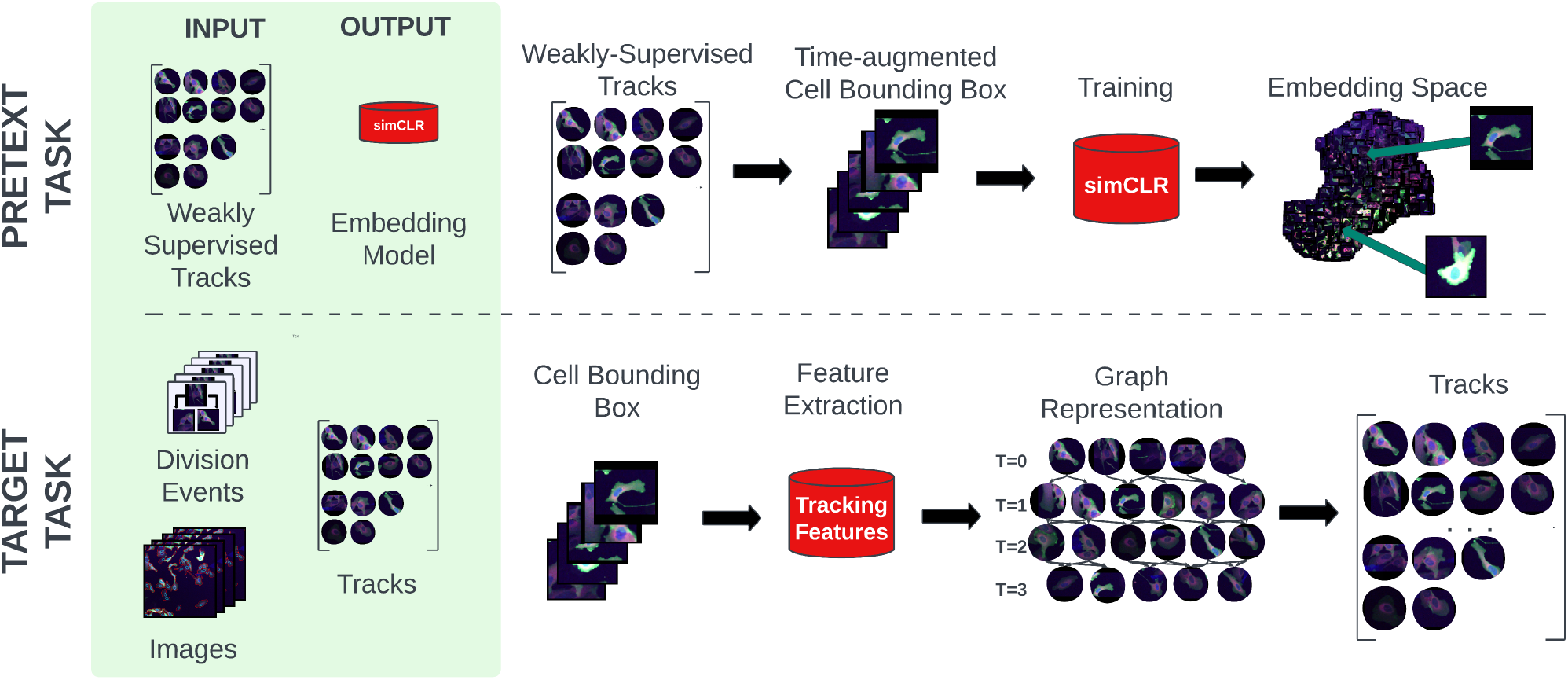
Tracking workflow: Our weakly-supervised tracking method consists of two main components: (1) The Pretext Task involves training a simCLR embedding model using time-based augmentations with weakly supervised tracks annotated by a segmentation-based tracker algorithm. (2) The Target Task utilizes features extracted from the simCLR embedding model applied without changes at inference in conjunction with additional hand-crafted and pretrained deep features, to optimize cell assignment over a graph structure and generate complete tracks.

In the next section, we will detail our contrastive learning approach, and in particular the variants we have developed for division detection and tracking. Then, we will detail our entire division detection and tracking methods, respectively.

### 3.3. Contrastive representation learning for cell division detection and tracking

#### 3.3.1. Contrastive learning approach

At the heart of our contrastive learning approach lies the simCLR model [8]. At its core, simCLR operates by generating multiple augmented versions of a given image and then training a neural network to identify which augmented views originate from the same base image. The key idea is to pull the representations of positive pairs (different augmentations of the same image) closer together while pushing the representations of negative pairs (augmentations of different images) apart. This is achieved through a contrastive loss function, known as the NT-Xent (Normalized Temperature-scaled Cross-Entropy Loss), which for a positive pair of examples is given by:

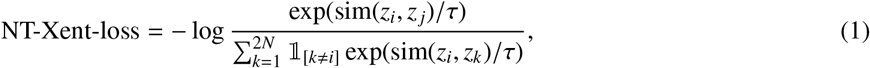

with *z*_*i*_ and *z*_*j*_ the representations of two augmented versions of the same image, sim(*z*_*i*_, *z*_*j*_) is the cosine similarity between these representations, *τ* the temperature scaling parameter, and *N* the number of images in the batch. The indicator function ⊮_[*k*≠*i*]_ equals 1 when *k* ≠ *i* and 0 otherwise. This formula essentially measures the probability that a pair of transformed images are recognized as originating from the same base image, normalized by the similarity of all other possible pairings in the batch.

In the context of weakly and self-supervised learning, one can categorize tasks into two types: pretext and target. Pretext tasks are solved during the initial training phase: the model learns representations from data by solving a task that is not ultimately useful but for which we have data available in large quantities. One can then use these learned representations to solve the target tasks, i.e. the one we actually want to solve but for which we have only limited expert-annotated data (e.g. specific predictive or classification tasks). In our case, these include identifying cell division events and tracking cell movements over time.

In this regard, the choice of the pretext task and the applied augmentations in the simCLR framework is paramount and can be seen as a form of supervision, as they directly influence the properties of the obtained representation space, which in turn influence its efficiency in helping to solve the downstream task [40]. For example, having rotations within the augmentations is useful to devise representations ultimately invariant to such rotations. In the following sections, we expose how we carefully craft augmentations to generate cell representations that ultimately help solving the tasks of cell division detection and tracking.

#### 3.3.2. Self-supervised contrastive learning for cell division detection

To generate cell representations useful for cell division detection, we train a simCLR model on a dataset of cell image samples, as illustrated in Figure 5(A). This dataset comprises 128 × 128 pixels patches associated with individually segmented cells. For each cell *c*, we consider its time-context as a triplet of cell patches 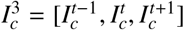, capturing information about both its immediate neighborhood and its motion. 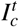 is a one zero-padded square patch centered around the cell located at a time *t*. Patches 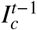 and 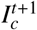 share the same spatial coordinates but situated in the preceding and the succeeding frame, respectively. Training the simCLR model with this variety of cell patches (and not only patches centered around the cells) ultimately proves useful to capture visual cues linked with cell division. For cells in the first and last frames of the time-series, the 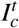 patch is repeated in place of the predecessor or successor patch. For instance, if a cell remains stationary, the patches at times *t* and *t*+1 would center around the cell. Conversely, if the cell undergoes significant movement, the patches at times *t* and *t* + 1 would reflect this spatial shift. A triplet 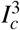 issued from 3 subsequent frames is illustrated in Figure 6.

**Figure 5.**
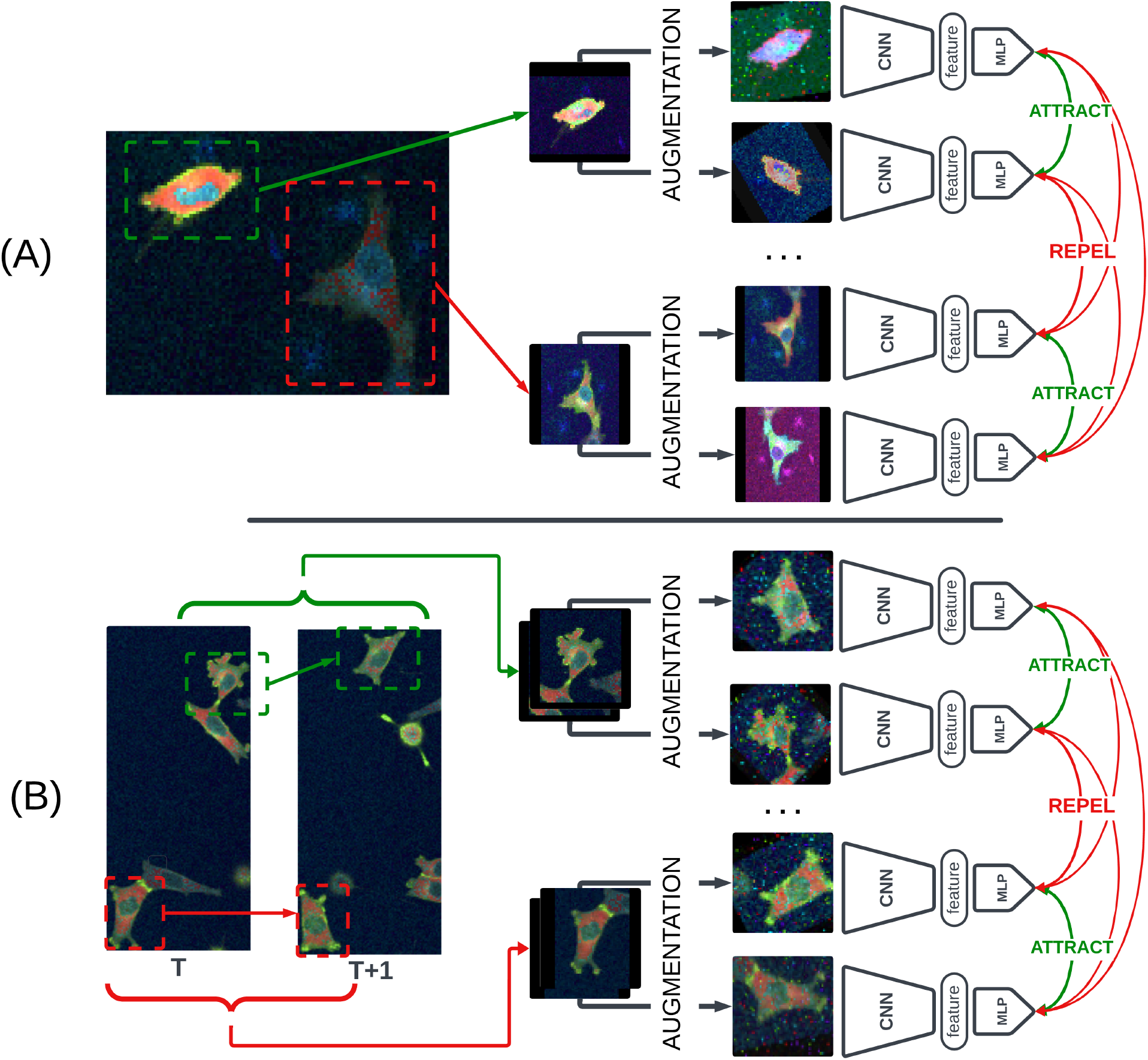
Division and tracking simCLR training diagram: In both these diagrams we represent the simCLR architecture as a Convolutional Neural Network (CNN) from which features are extracted as well as a Multi-Layer Perceptron (MLP) used for the training only. **(A)** Division-simCLR training: For each individual cell in the training videos, we extract the image patch corresponding to the cell’s Region of Interest (ROI) and scale it to a constant size. At training, for each sample patch, we apply our stochastic augmentations, propagate the individual images through the model and subsequently train the model with the objective of matching augmented images issued from the same sample and repelling them in the embedding space from the other samples’ augmented images. **(B)** Tracking-simCLR training: Given two frames with weakly supervised tracking labels, for each pair of corresponding cells within a track, we extract image patches corresponding to the cell’s ROIs in their respective frames. At training, we apply stochastic augmentations on both patches and propagate them through the model. We train the model with the objective of matching image patches issued from the same track and to repel them from other pairs in the embedding space.

**Figure 6.**
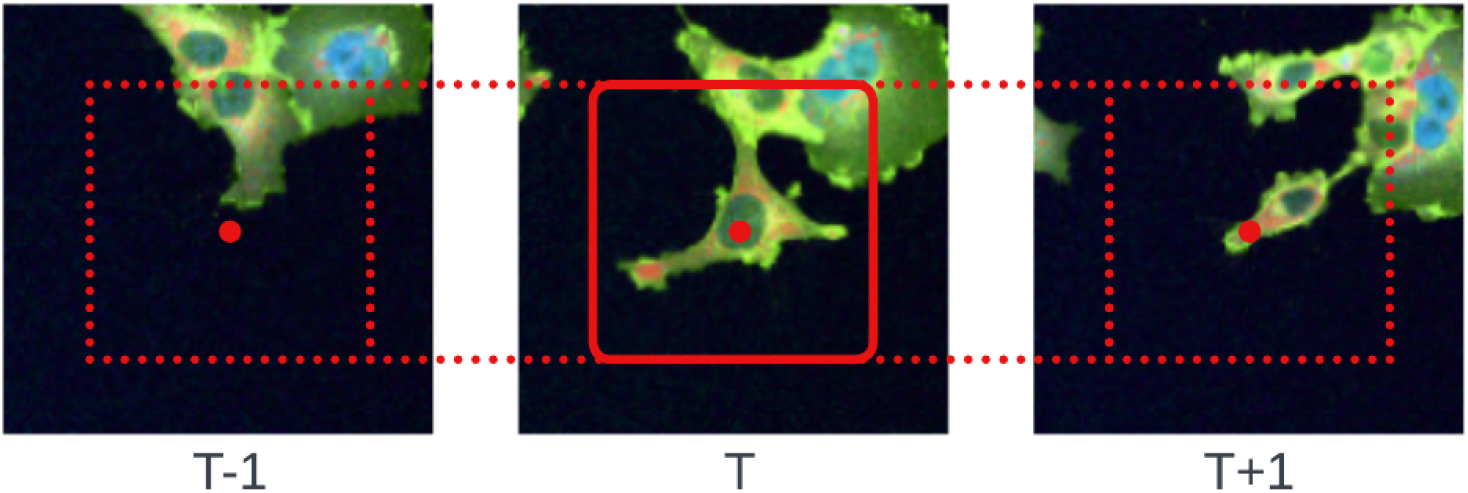
Temporal context extraction process for a cell at time T. Starting with the cell and its segmentation, a padded square patch is extracted from its frame using its Region of Interest (ROI). The time-context information incorporates the ROIs at corresponding coordinates in the preceding T-1 and following T+1 frames. The notation 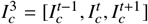 denotes the time-context image patches for a cell *c*.

In the training of the simCLR model, each image within a triplet 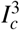 is treated independently. We duplicate the image dataset and apply distinct augmentations to each individual image. These augmentations are illustrated in Fig. 7. We use augmentations from the original simCLR paper as well as augmentations linked to fluorescence microscopy: (i) Poisson noise to emulate the noise generated by the fluorescence microscope imaging [41] [42], (ii) Additive White Gaussian Noise (AWGN) to emulate the variation in the expression of the lit-up fluorescent pixels and random background noise [43], (iii) Salt and Pepper Noise to emulate a variation in the location and expression of the fluorescent reporters by simulating activation or inhibitions of fluorescent proteins[44]. Finally and importantly, we introduce a novel augmentation technique that we name channel switching, whichconsists in randomly permuting fluorescent channels. This augmentation can be interpreted as a substantial color augmentation while also diminishing the model’s reliance on the expression of individual fluorescent reporters, resulting in more robust and morphology-oriented features. We denote those features *division-simCLR*. The distance between features is calculated using the cosine distance with which the simCLR model is trained. To characterize a triplet 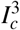, we can extract 3 division-simCLR features and stack them, which we denote *division-simCLR*^3^.

**Figure 7.**
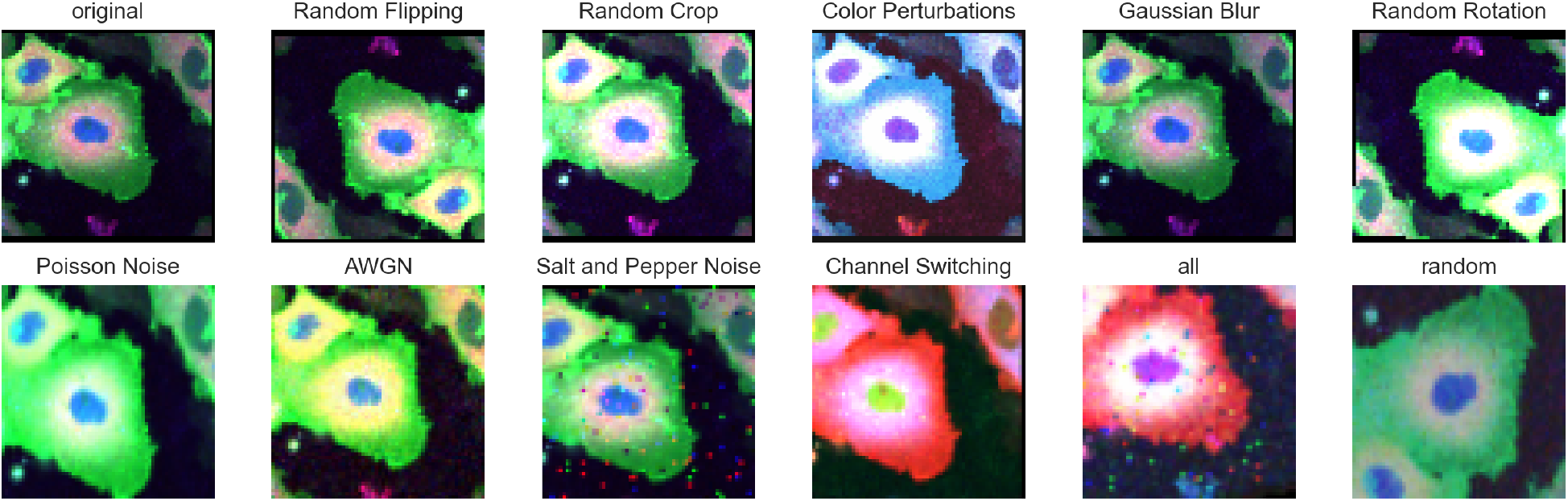
Augmentations employed during the training of our division-simCLR model. The original augmentation methods from the simCLR paper (flipping, cropping, color perturbations, Gaussian blur, rotation) are included, alongside microscopy-specific augmentations (Poisson noise, Additive White Gaussian Noise, and Salt and Pepper noise). Additionally, channel switching is introduced as the process of randomly permuting image channels in order to prioritize the impact of morphology over fluorescent proteins and their spatial distribution in the resulting feature space.

#### 3.3.3. Weakly-supervised contrastive learning for cell tracking

To generate cell representations useful for cell tracking, we propose a novel weakly-supervised constrastive learning approach. It consists in using an unsupervised (although sub-optimal) tracking algorithm to generate cell tracks from video-microscopy datasets, and then train a simCLR model to generate representations such that consecutive cells in tracks are close to one-another in the resulting representation space. This approach is illustrated in Figure 5(B).

We use Lineage Mapper [7] as unsupervised tracking algorithm to generate a weakly-supervised dataset of pairs of consecutive cells (i.e. cell matches). It uses auxiliary information (not directly contained within the images) about cells to generate tracks, namely the cell’s spatial distance, overlap and size ratio (computed from cell segmentations). While the resulting set of tracks does not attain the same quality as human annotations or supervised tracking approaches, it can be amassed at scale without requiring human intervention. Validation upon annotated tracks confirms that this method achieves a Tracking Fraction (TF) score of 0.75, representing the average correctness of links within tracks (as defined later in section 3.6.3). This level of performance enables the representation model to acquire generalized embeddings pertinent to the tracking task. The resulting dataset comprises pairs of cell matches, each patch being of size 128 × 128 pixels and centered around the cell.

Subsequently, we train a simCLR model to generate representations of cell matches based on the weakly-supervised dataset of cell tracks. However, instead of training the model to match different augmentations of the same image, we adapt the approach to match the same cell at two consecutive time-points from within the tracks. This adaptation effectively incorporates time-shifting as an augmentation technique within the standard simCLR framework, allowing us to learn cell representations that are optimized to remain similar over time. Furthermore, we apply classical simCLR augmentations along with microscopy-specific augmentations as described in section 3.3.2. We denote the resulting features *tracking-simCLR*, and use the cosine distance to compute the distance between two cell representations.

### 3.4. Cell division detection with contrastive features

Our cell division detection method operates as a two-component framework, as previously explained in section 3.2. It is illustrated in Figure 3: firstly, a self-supervised pretext task for the purpose of generating representations, and secondly, a supervised target task designed for division detection. This way, the division-simCLR^3^ representations generated by the self-supervised task are harnessed within the supervised training of a cell division detection model. This framework is represented in Figure 3. In this section, we explain how we use the division-simCLR^3^ features derived from the pretext task in the context of the division detection target task. We first introduce the graph structure and associated features we use to model candidate cell triplets as possible mother and two daughters, and then explain how we classify such triplets using a dual-input classifier specifically tailored for identifying division events.

#### 3.4.1. Graph modeling and features for cell division detection

We initially select candidate cell triplets from the video, i.e. a possible mother cell at frame *t* and associated two daughter cells at frame *t* + 1. These triplets are then classified as either positive or negative division events. This classification problem is highly imbalanced as the number of division events is very low compared with the number of potential mother-daughters candidate in a video. We generate a graph representation of the cell population in the video over time to select candidate cell triplets and match potential mother cells with their potential daughters. For a video spanning *T* frames, where each frame *I*^*t*^ contains *N*^*t*^ cells 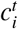, we construct a graph 𝒢 = (𝒱, *ℰ*) with a vertex set 𝒱 and an edge set ℰ. We create a vertex 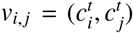 for every pair of cells within the same frame (except the first frame) which centroids are distant from less than *D*_max_. Subsequently, for each pair of cells in 𝒱, we search the preceding frame for cells that maintain a mean centroid distance to the cells 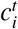 and 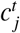 smaller than *D*_max_. If a cell 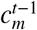 satisfies this criterion, it is incorporated into the vertex set as 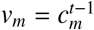, and an edge *e*_*m,i, j*_ = (*v*_*m*_, *v*_*i, j*_) is created. Thus, each edge in ℰ represents a mother-daughter candidate triplet. We select *D*_max_ to match 95% of our annotated division events and not 100% so as to remove (and deliberately ignore) outlier triplets. Reducing the number of candidates selected overall is key to not only make sure we consider as many true positives as possible, but that we filter out as much negatives as possible to reduce the class imbalance in the classification dataset.

We then derive cell representations to be used to solve the cell division classification task. For every edge in the graph representing a potential division event triplet, we compute the time-context division-simCLR^3^ representations for the mother cell and its two potential daughter cells using the model described in section 3.3.2. Given a candidate mother cell *c*_*m*_ alongside its two prospective daughters 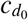 and 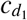, we obtain triplet image patches 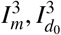, and 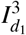, centered in the respective cells’ centroid. These image patches yield division-simCLR representations denoted by division-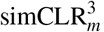, division-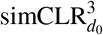, division-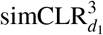, collectively forming nine vectors each of size 2048. For each potential division event represented by an edge in the graph, we additionally calculate a set of hand-crafted features to evaluate the candidacy of a mother cell *c*_*m*_ and its two potential daughter cells 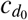 and 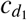. These handcrafted features include:

*Spatial Euclidean distances* between the mother candidate and each daughter candidate, as well as between the two daughter candidates: 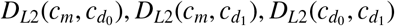.

- *Size distance* between the daughter candidates, based on their areas (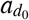 and 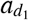), calculated as: 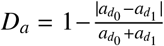.
- *Aspect ratio* distance between the daughter candidates, using their aspect ratios 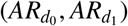, computed as: 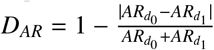.
- *Pairwise cosine distances* within the division-simCLR embedding space for the candidates *c*_*m*_, 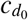, and 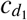 with their embeddings division-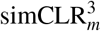, division-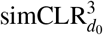, division-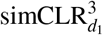, by calculating the cosine distance between each pair division-simCLR_*i*_, division-simCLR _*j*_ of the 9 total embeddings as: 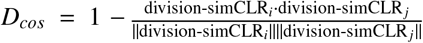. This results in 36 unique pairwise cosine distances for the 9 embeddings.

Thus, for any edge 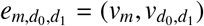 in ℰ, we construct a handcrafted feature vector 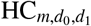 encapsulating all 41 distances.

#### 3.4.2. Cell division detection classifier

Once equipped with candidate cell triplets representations and associated graph structure as introduced in section 3.4.1, we can use them within a dual-input classifier specifically tailored for identifying division events. It processes both the division-simCLR^3^ embeddings and hand-crafted feature vectors simultaneously to perform its classification task. The architecture of our model is made of two main components, as illustrated in Figure 8: (1) a recurrent encoder that condenses the dimensionality of the embeddings, where the input consists of concatenated cell triplets’ division-simCLR embeddings along the time dimension, capturing the temporal relationship between mother and daughter samples, and (2) a binary classifier that integrates the recurrent encoder’s hidden state with the hand-crafted feature vector to make its classification. The recurrent encoder employs an Long Short-Term Memory (LSTM) network with a hidden layer size of 64 across two layers, including a dropout rate of 0.2 to prevent overfitting. The binary classifier Multi-Layer Perceptron (MLP) utilizes four linear layers with intermediary sizes of 64, 32, 16, and 8, activated by ReLU functions, leading to a final classification layer activated by a sigmoid function.

**Figure 8.**
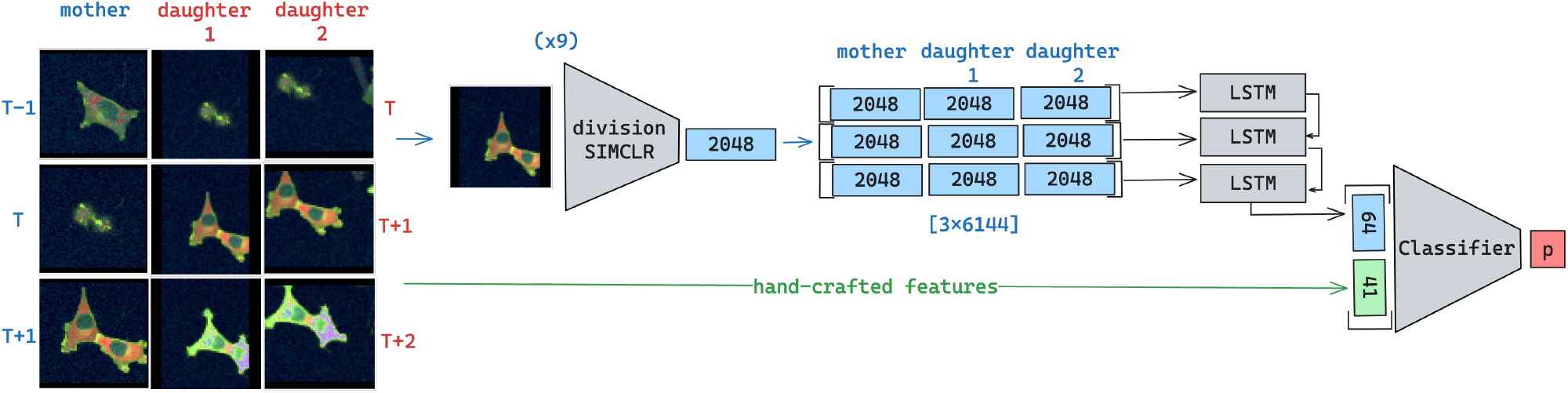
Diagram illustrating the architecture of our division detection classifier. The classifier utilizes division-simCLR^3^ features and hand-crafted features via two sequential models: (1) a recurrent classifier (LSTM), where the input consists of concatenated cell triplets’ division-simCLR^3^ embeddings along the time dimension. The LSTM generates a latent state, which is subsequently concatenated with the hand-crafted features. (2) The resulting vector is then fed into a binary classifier (MLP), which outputs the class probability of the division event.

The model is trained using a weighted binary cross-entropy loss and optimized with ADAM gradient descent [45], with sample weights computed to balance the dataset’s class imbalance among candidate division triplets. We adopt a learning rate of 1e−5, which is gradually adjusted using cosine annealing over a period of 50 epochs to a minimum learning rate of 1e−7. To address dataset imbalances and avoid bias from any specific annotated video - and thus its experimental conditions - we upsample examples from videos with fewer division events to equalize the number of division events across the training dataset. Furthermore, we use aggregated ensemble learning. With *k* the total number of videos used for the training and *p* the number of videos used for training of each fold. For each fold, we use (*k* − *p*) videos for training and evaluate on the remaining *p* unseen videos. For each fold, the (*k* − *p*) training-split videos are themselves divided into a training set (80 %) and a validation set (20 %). The latter is used for early stopping. This approach results in *k/p* models, each trained on (*k* − *p*) training videos. During inference, we aggregate predictions from all models, averaging the probabilities for assignment to the graph’s edge.

Finally, at inference, once all graph edges are classified we ensure division consistency by iteratively choosing edges with the highest probability scores and removing their corresponding nodes until no positively classified edge remains. We are thus able to identify mother-daughterstriplet associated with division events, using only minimal annotations for the classifier by leveraging the self-supervised contrastive features.

### 3.5. Cell tracking with contrastive features

In this section, we present our methodology for cell tracking in scenarios characterized by low temporal resolution. First, we elaborate on the set of features employed in our tracking method, including those generated through weakly supervised contrastive learning as presented in section 3.3.3. Second, we expose how we model cells over time using a graph for tracking. Finally, we present our graph assignment technique named Sliding Window Assignment Graph (SWAG), which effectively leverages these features and graph modeling to extract tracks from time-series of cell images.

#### 3.5.1. Tracking features

Our cell tracking methodology integrates a wide array of features including spatial, hand-crafted, and deep learning-based features. Each feature type, paired with a specific distance metric, contributes to a comprehensive composite distance metric facilitating nuanced comparisons between cell pairs.

First, we use spatial features that are traditionally used in tracking applications [7]:

- *Centroid distance*: For two cells 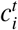 and 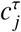 we compute the Euclidean distance *D*_centroid_ between their centroids as measure of their spatial separation.
- *Overlap Metric*: The overlap-based distance metric *D*_overlap_ assesses the degree of spatial intersection between two cells 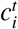 and 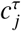. It is defined as 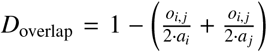, where *a*_*i*_ and *a*_*j*_ represent the pixel areas of the respective cells and *o*_*i, j*_ their intersection.
- *Size distance: We calculate the size difference between two cells,* 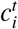 *and* 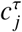, *in terms of their pixel areas, a*_*i*_ *and a*_*j*_. The distance metric *D*_size_ reflects the relative size variation. It is defined as 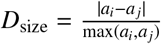.

We investigate several deep learning feature sets with respect to their performance in our framework from both standard pre-trained classification models and custom-trained models:

- *simCLR features:* We use the features introduced in section 3.3.2, namely division-simCLR features to characterize individual cells and division-simCLR^3^ features to characterize the temporal context 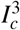 around each cell *c*. When using division-simCLR^3^ features, the distances between cell pairs are determined by synchronizing the embeddings with respect to their temporal stamps.
- *Weakly-supervised contrastive learning features*:We use the tracking-simCLR introduced in section 3.3.3. Those features are optimized for the tracking of cells using weakly supervised contrastive learning.
- *Pretrained deep features:* We use the ResNet-50 [46] and the AlexNet [47] models to compute embeddings (internal representations just before the classification layer) from 128 × 128 bounding boxes centered around individual cells. This process yields a 2048-dimensional vector for ResNet-50 and a 4096-dimensional vector for AlexNet, that we denote *resnet* and *alexnet*. When considering two cells 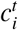 and 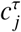, we extract their respective representation embeddings e.g. 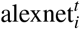 and 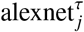, and compute the distance between these cells using the L2 norm: 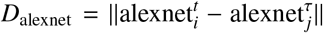. Furthermore, we use the time-context feature extraction technique introduced in Section 3.3.2 where we embed the image triplet 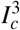 for each cell. To quantify the distance between two cells using this approach, we sum the L2 norms of their temporally aligned embeddings. These features are referred to as *alexnet*^3^ and *resnet*^3^, depending on the model employed.

Importantly, our methodology is not confined to specific features or distance metrics. It is designed as a flexible framework, adaptable to various feature types suitable for the dataset at hand. Table 3.5.1 summarizes the various features used in our tracking methodology, along with their corresponding distance metrics:

#### 3.5.2. Graph modeling for tracking

Our approach to cell tracking uses a graph structure to represent cells over time and capitalize on the features described in section 3.5.1 to characterize individual cells. Cells are vertices in the graph, and cell linkages are influenced by constraints such as time, localization and division events. These constraints determine the edges that exist within the graph. The edge weights capture the similarity between cells in the feature space. By identifying optimal paths within this graph, we can delineate cell tracks throughout the video that strictly adhere to predefined conditions, e.g. a cell without a successor marks the end of a track.

To model cellular movements within a video we construct a weighted directed graph denoted as 𝒢 = (𝒱, *ℰ*), with 𝒱 the set of vertices corresponding to individual cells, and ℰ the edges indicating potential connections between these cells. Each vertex *v* in 𝒱 represents a cell object within an image frame, identified as 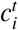, where *i* is the cell’s index in the frame, and *t* is the frame’s relative timestamp. The edges in ℰ connect vertices from 𝒱 based on specific criteria:

1. An edge *e* in ℰ can only connect cell objects from one frame to the next, i.e. between times *t* and *τ* = *t* + 1.
2. A vertex *v* modeling a cell 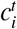 which is a mother in a division event cannot have outgoing edges.
3. Similarly, a vertex *v* modeling a cell 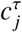 which is a daughter in a division event cannot have ingoing edges.
4. An edge is established between two vertices *v*_*i*_ and *v*_*j*_, corresponding to cells 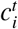 and 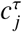, if the spatial distance between them, calculated as 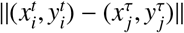, is below the maximum distance threshold *D*_*max*_.

The weights of edges in ℰ are determined by composite distances derived from features extracted from the associated cell objects. For a pair of vertices *v*_*i*_ and *v*_*j*_, representing cells 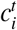 and 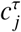, the edge weight *w*_*i j*_ is calculated using the following formula:

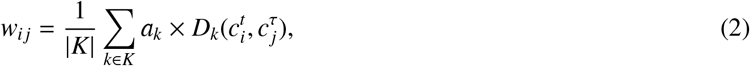

where *K* is the set of features related to the cell objects, *D*_*k*_ is the distance function applied to these features, and *a*_*k*_ represents the weight assigned to each feature’s distance. The features are normalized to have a mean of 0 and a standard deviation of 1 over the entire video.

#### 3.5.3. Sliding Window Assignment Graph (SWAG)

In this section, we introduce an innovative graph optimization method named Sliding Window Assignment Graph (SWAG). The SWAG method enables to extract tracks from the graph constructed as explained in section 3.5.2. The main idea is to assess cell linkage under varying multiple initial conditions to enable a robust track extraction. SWAG allows for a probabilistic rather than a deterministic track evaluation, which enables several evaluation of each cell’s role within a track but also allows for the consideration of various track dynamics influencing cell matching within the reference of the sub-graphs. Hereafter we detail the different steps of the SWAG approach that encompasses extracting sub-graphs, generating tracklets from these sub-graphs, and ultimately compiling global tracks across the entire video by leveraging these tracklets. The workflow of the SWAG method is illustrated in Figure 9.

**Figure 9.**
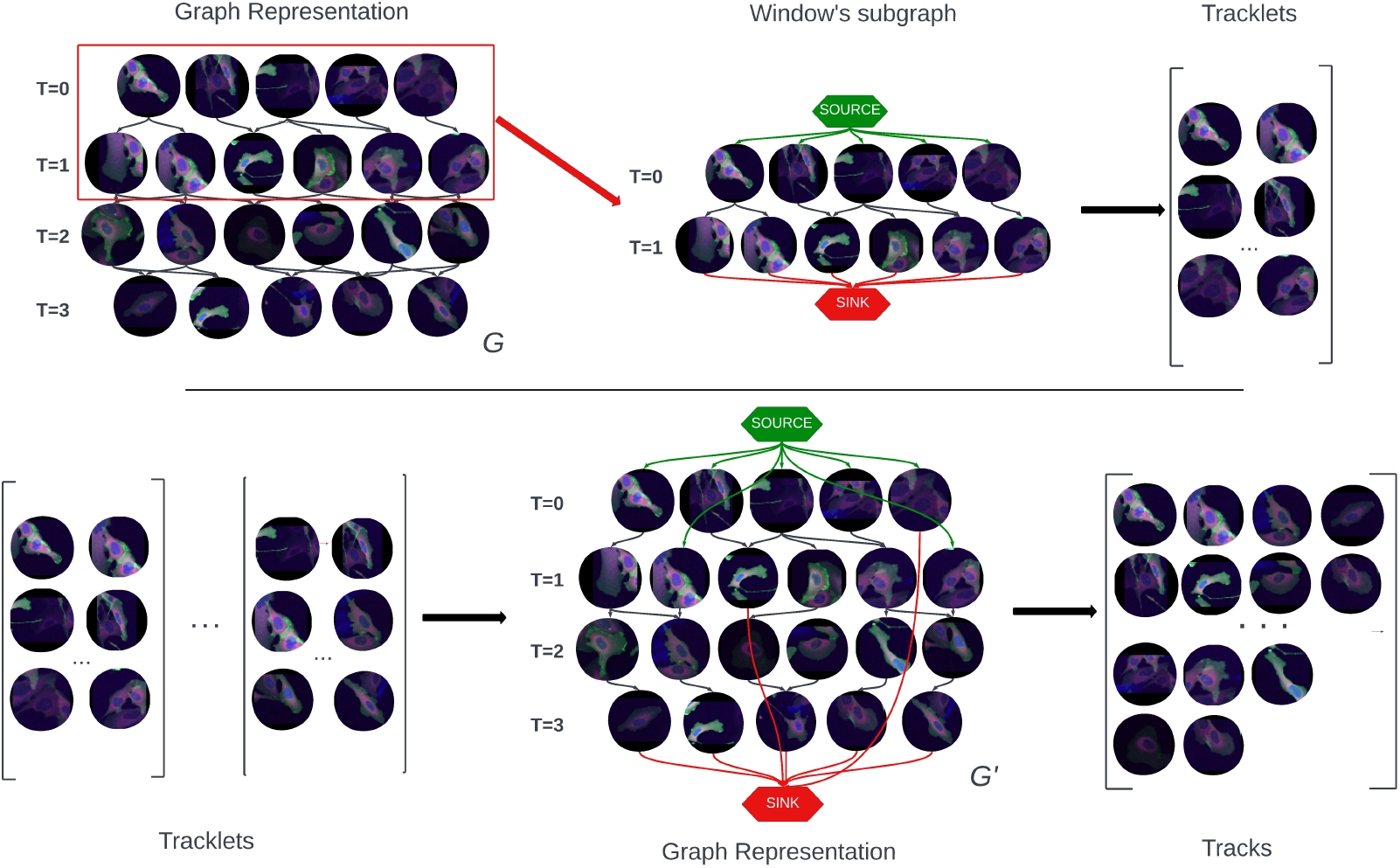
Diagram illustrating the Sliding Window Assignment Graph (SWAG) Method. **Top:** We first build a graph 𝒢 containing all cells in the video, connected from one frame to the next with the application of constraints (based on localization and division events). We then iteratively extract sub-graphs from it using a rolling window. For each sub-graph, we compute the shortest path using edge weights computed on cell’s features. Each shortest path extracted from each sub-graphs is collected as a Tracklet. **Bottom:** Using the collected tracklets we build a new graph 𝒢^′^ with vertices for every cell in the video and edges only for links showing in the tracklets set. Each edge is weighted based on the number of occurences of its link in the set of tracklets. We then use a Markov Decision Process with policy iteration to extract final complete tracks from 𝒢^′^.

To extract tracks out of the initial graph, we implement a rolling-window method to facilitate a structured and detailed examination of the temporal dynamics and lineage relationships of cells throughout the time series, all the while allowing each cell to be considered at least once as the potential start or end of a track. This approach, involving the definition and analysis of rolling window sub-graphs, is characterized by two parameters: the window size *l* and the stride *s*. Using a rolling window, we can thus define a sub-graph 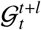 as the subset of the larger graph 𝒢 that includes only the nodes corresponding to cell objects in frames from timestamp *t* to *t* + *l*. We systematically extract such sub-graphs from 𝒢, beginning at *t* = 0 and incrementing *t* by *s* until we cover the entire duration of the image time series, denoted by *T*. Furthermore, in each of these sub-graphs, 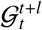, we introduce two new vertices, *v*_*S*_ and *v*_*T*_, which serve as the source and sink, respectively. These two vertices serve as placeholders for the beginning and end of tracks: A track in the graph can be any path between *v*_*S*_ and *v*_*T*_. We connect these vertices to the existing vertices in the sub-graph using directional edges weighted at 0. We create an edge from *v*_*S*_ to a vertex *v*_*i*_ if *v*_*i*_ has no incoming edges, indicating the absence of predecessors for *v*_*i*_ within the sub-graph 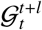. Conversely, we establish an edge from a vertex *v*_*j*_ to *v*_*T*_ if *v*_*j*_ lacks outgoing edges, implying no successors for *v*_*j*_ within the same sub-graph.

We then compute in each sub-graph a set of tracklets using a method that involves the following steps:

1. We calculate the shortest average path between the source node *v*_*S*_ and the sink node *v*_*T*_ within the sub-graph 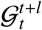, using a procedure described in Algorithm 1 below. This calculation defines a tracklet, denoted as 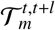, which includes all vertices present along the identified path.
2. Next, we remove the vertices that are part of the tracklet 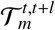 from the sub-graph, effectively updating the graph’s structure.
3. The process is repeated: if the sub-graph still contains vertices other than *v*_*S*_ and *v*_*T*_, we return to step 1 and continue the computation. The process terminates when only *v*_*S*_ and *v*_*T*_ remain in the sub-graph.

##### Algorithm 1 Algorithm for Shortest Average Path

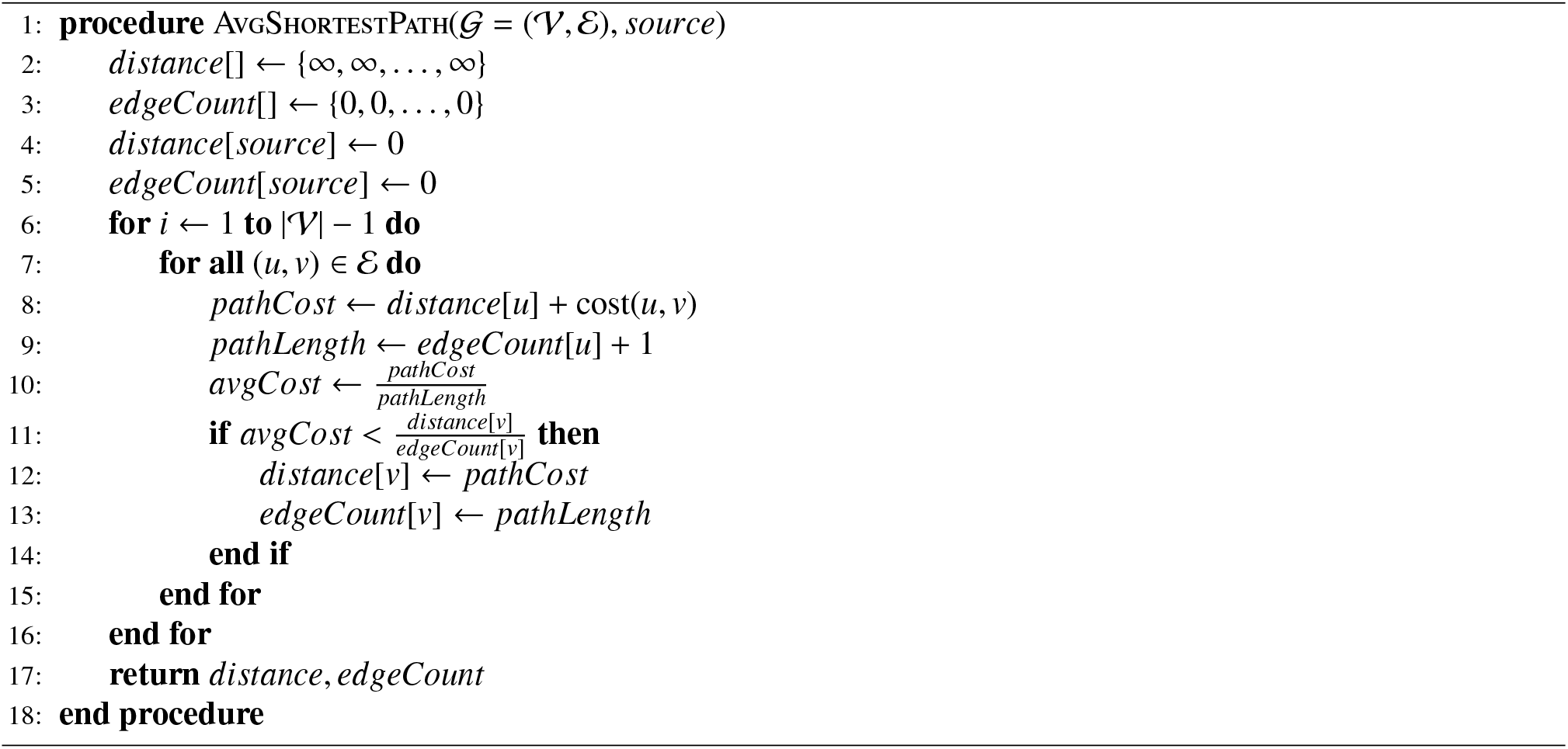

We then use the tracklets to generate transition probabilities between nodes in the graph. The tracklets we have extracted indeed represent possible linkage between cells in multiple initial video conditions (duration, start and end). After computing tracklets for all sub-graphs, we create a comprehensive set of tracklets that spans the entire graph 𝒢. This set enables us to determine the frequency of selection for each edge between two nodes in the graph. Here, the edge frequency indicates how often a link between two nodes appears across the entire set of tracklets. Using this data, we construct a new directed graph 𝒢^′^ = (𝒱^′^, *ℰ*^′^). The vertex set 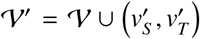 supplements the original graph vertices by the addition of source and sink vertices 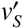 and 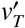. Thus, these vertices correspond to cells 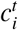 and 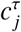 or represent the start and terminal states of a tracklet. Then an edge 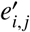 in ℰ^′^ connects two vertices 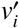 and 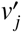 if and only if there is at least one connection between these vertices in the set of previously computed tracklets. We then transform the occurence of each edge 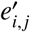 in the set of tracklets into a transition probability, represented as *p*_*i, j*_. This is defined as:

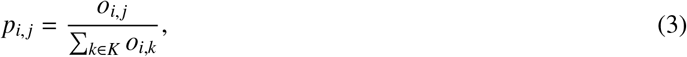

where *K* denotes the set of successor vertices from the node *v*_*i*_, and *o*_*i, j*_ refers to the frequency of the edge 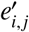 linking vertices 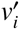 and 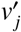. in the set of tracklets This calculation of transition probabilities provides a quantitative basis for determining the likelihood of cell transitions within the graph, grounded in the observed occurrences within the tracklet dataset.

Finally, once provided with this probabilistic graph 𝒢^′^, we can use a Markov Decision Process (MDP) to extract the final tracks. This MDP has the following parameters:

- The set of states, denoted as *S*, is defined as 𝒮 = 𝒱^′^.
- The set of available actions in a given state *s*, represented as 𝒜_*s*_, is defined as 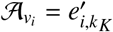, where *K* is the set of successor vertices of 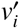 in 𝒢^′^.
- The transition probability, *p*(*s, s*^′^), quantifies the likelihood that action *a* in state *s* at time *t* leads to state *s*^′^ at time *t* + 1. Formally, this is expressed as 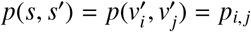.
- The immediate reward for transitioning from state *s* to state *s*^′^ due to action *a, R*_*a*_(*s, s*^′^), is calculated as 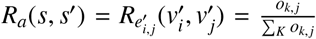, where *K* represents the set of predecessor vertices of 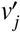. This reward model is applicable to all transitions, except for the transition to the terminal state 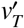, where the reward is set to zero.
- The policy function *π* maps actions from 𝒜 to states in 𝒮.

To optimize the policy *π*, we apply the policy iteration algorithm [48]. This involves iteratively calculating a value function over the state space using the formula:

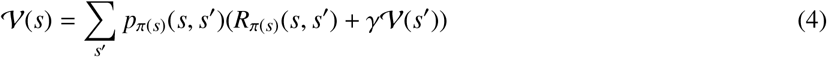

Here, *γ* is the reward horizon parameter, set at 0.9. The policy is then optimized as follows:

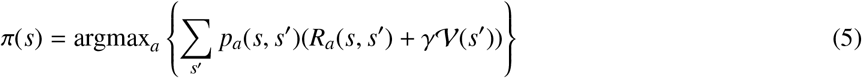

This iterative process continues until convergence is achieved, yielding the optimal policy *π*^*^. To extract a track from 𝒢^′^, we initiate at the start vertex *v*_*S*_, following the optimal policy *π*^*^ to the terminal vertex *v*_*T*_. Vertices along the track are then removed from 𝒢^′^, and this process repeats until only the start and terminal nodes remain in the graph. This method effectively returns a set of tracks, each denoted 𝒯_*m*_, such that 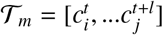 for a track of length.

Furthermore, we introduce an extension to the SWAG method presented above that enhances the extraction of tracklets by varying the size of the sliding window, that we name Pyramidal SWAG, or P-SWAG. P-SWAG consists in executing the sliding window technique multiple times with different window sizes and aggregating the resultant tracklets from sub-graphs of these varying sizes, thus we create creating an augmented graph 𝒢^′^ through a pyramidal strategy. This approach enables the consideration of a broader array of sub-graph states, each of a different size, thereby influencing the transition probabilities between cells in the Markov Decision Process (MDP).

### 3.6. Evaluation metrics

This section outlines the metrics employed to evaluate cell division detection, the quality of cell tracking features, and cell tracking performance.

#### 3.6.1. Cell division detection evaluation metrics

To assess our cell division classifier, we employ the F1-score, a harmonic mean of precision and recall, alongside the area under the precision-recall curve (AUC), and the model’s Accuracy. These measures provide insights into the model’s performance, especially in dealing with class imbalances, with scores ranging from 0.0 to 1.0—where scores closer to 1.0 indicate better performance.

In the context of learning outcomes, we calculate these metrics through cross-validation, ensuring a unified evaluation across all test folds for a direct comparison of trainable and non-trainable models. This approach aggregates predictions to produce a single, comprehensive performance score.

#### 3.6.2. Cell tracking features evaluation metrics

To evaluate our tracking features, we utilize a set of metrics to precisely measure the accuracy of our tracking algorithm. These metrics enable an in-depth analysis of how individual features or their combinations perform in correctly matching cells across frames.

***Top-1 (Whole Frame)*** measures the accuracy of the algorithm in identifying the best tracking match in the subsequent frame, highlighting the precision of the tracking feature. ***Top-3 (Whole Frame)*** expands this assessment to whether the correct match falls within the top three selections in the next frame. Both ***Top-1 (In Range)*** and ***Top-3 (In Range)*** limit this evaluation to cells within a 100-pixel radius, focusing on the performance of the feature in tracking spatially proximal cells. Scores for these metrics range from 0.0 to 1.0, where a score closer to 1.0 denotes effective tracking capability.

***Mean Rank (Whole Frame***/***In Range)*** calculates the average rank of the correct match, either globally or within the specified proximity, offering an overarching view of the feature’s tracking accuracy. A lower mean rank indicates higher efficiency in identifying correct tracking matches.

#### 3.6.3. Cell tracking evaluation metrics

We evaluate cell tracking methodologies using established metrics from cell tracking research, ensuring a thorough assessment of tracking performance in cell screening applications.

***Track Reconstruction Accuracy (TRA)***, following Matula et al.’s [49] methodology, uses the normalized Acyclic Oriented Graph Matching (AOGM) measure for evaluating the precision of tracking algorithms in reconstructing cell trajectories. This metric serves as a benchmark metric for understanding how accurately an algorithm follows cell paths over time, although it is not the most interpretable metric especially in the context of biocomputing applications.

***Complete Tracks (CT)*** assesses the percentage of cell trajectories fully reconstructed by the tracking algorithm, with any error disqualifying the entire track.

***Track Fraction (TF)*** evaluates the average completeness of the reconstructed tracks, providing an aggregate measure of tracking success across the dataset.

The ***Partially Reconstructed at 70% (PR@0.7)*** metric, meanwhile, considers the portion of cell tracks reconstructed to at least 70% completeness, offering insights into the algorithm’s capability to accurately follow significant segments of cell trajectories.

Scores for all these metrics fall between 0 and 1, where a score of 1 represents optimal tracking performance.

## 4. Results

### 4.1. Cell division

#### 4.1.1. Deep feature analysis

We compare the division-simCLR features introduced in Section 3.3.2 with pretrained deep features alexnet and resnet based on the AlexNet and ResNet backends, as introduced and explained in Section 3.5.1, in the paragraph “Pretrained deep features”.

For each of those base features, we consider two types of representations: (i) one using a temporal-context, denoted as 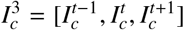 and leading to features division-simCLR^3^, alexnet^3^ and resnet^3^; and (ii) the other using a single-frame representation, 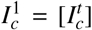 leading to features division-simCLR, alexnet and resnet. The results of this analysis are presented in Table 2. They are computed using leave-p-video-out cross-validation, with 5 folds each with testing set size of 2 videos. We iteratively consider 20% of our annotated dataset as test data and train our model on the remaining data, until the entire dataset has been evaluated.

**Table 1.**
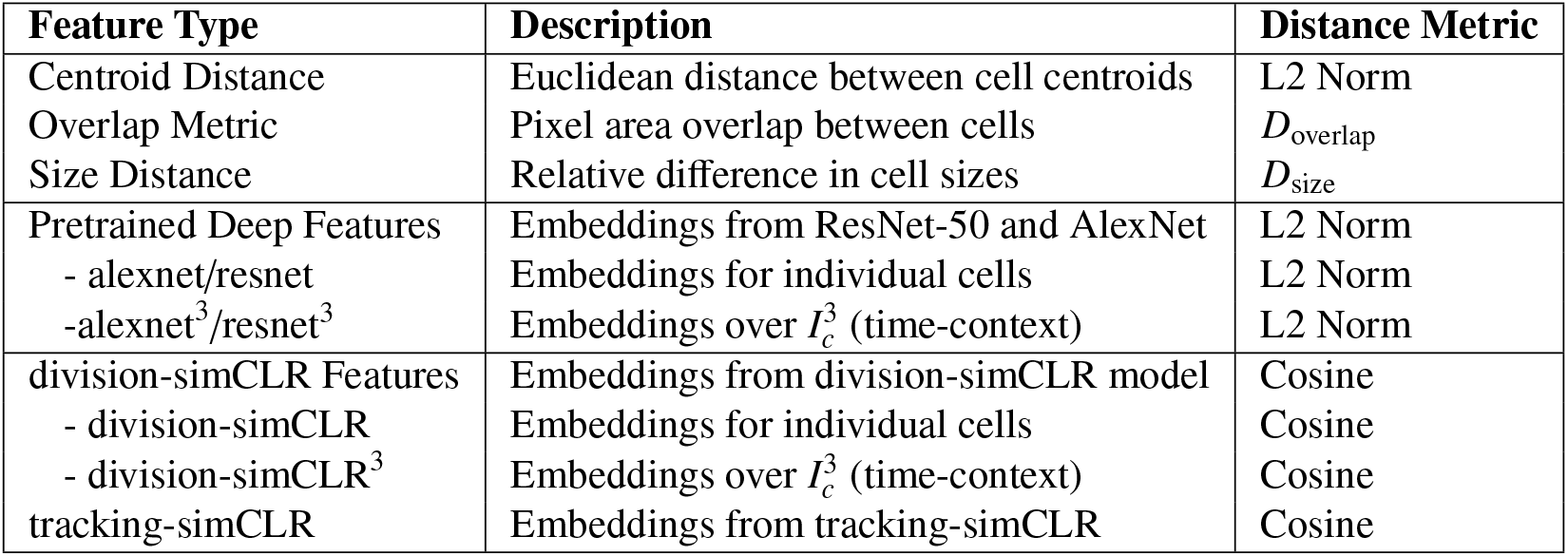
Summary of features and associated distance metrics used in our tracking methods.

**Table 2.** Evaluation of the performance of the division detection method with and without our contrastive features and the inclusion of the time-context. We assess these variations of our tracking algorithm on our dataset (A375 with a temporal resolution of 240min) using the AUC, the F1-score, the Precision, the Recall and the Accuracy.

| Method | Feature Type | AUC | F1 | Precision | Recall | Accuracy |
| --- | --- | --- | --- | --- | --- | --- |
| **Single Time-Point** | **resnet** | 0.1070 | 0.1931 | 0.1878 | 0.2189 | 0.8978 |
|  | **alexnet** | 0.1193 | 0.3041 | 0.3491 | 0.2754 | 0.9310 |
|  | **division-simCLR** | 0.2931 | 0.4263 | 0.4778 | 0.3957 | 0.9442 |
| **Time-Context** | **resnet<sup>3</sup>** | 0.1434 | 0.3686 | 0.3997 | 0.3870 | 0.92476 |
|  | **alexnet<sup>3</sup>** | 0.1955 | 0.3188 | 0.3081 | 0.3315 | 0.92237 |
|  | **division-simCLR<sup>3</sup>** | **0.3641** | **0.4692** | **0.6324** | **0.4018** | **0.9550** |

Our results indicate that pretrained deep features achieve an F1-score of up to 0.36 when considering the time-context and using resnet^3^ features. In contrast, relying solely on single time-point data decreases performance, with the F1-score dropping to a maximum of 0.30 when using alexnet features. Using simCLR features as introduced in this work significantly improves outcomes, reaching an F1-score of 0.42with division-simCLR features and rising to 0.48 for division-simCLR^3^ features with the addition of the time-context. Correspondingly, the Precision in division detection with division-simCLR^3^ features increases from 0.47 to 0.63 with the inclusion of time-context. These findings affirm the efficacy of our proposed approach, both regarding the use of contrastive features as well as the integration of the time-context.

#### 4.1.2. Cell division detection method benchmarking

In this segment of our study, we conduct a comparative analysis of our method against the Lineage Mapper’s (LM) division detection algorithm [7]. This comparison is relevant as both LM and our method can be used for semantic division detection and require few annotations.

We undertake this comparative assessment across four distinct datasets: our A375 cell line dataset and three datasets from the Cell Tracking Challenge (CTC) that presented a sufficient number of division events for effective training of our methodology. It is important to note that the CTC dataset results were achieved using alexnet^3^ features instead of division-simCLR^3^ features, due to constraints related to the size of these datasets, as well as their diversity w.r.t. cell lines and imaging parameters.

The outcomes of this benchmarking exercise are detailed in Table 3. The results underscore the performance of our method across both the A375 and CTC datasets. We prioritize precision over recall to minimize the impact of false positives and ensure a high likelihood of correct division detection, thus favoring the usefulness of the tracking system for downstream analysis.

**Table 3.** Benchmarking of our proposed division detection method against the Lineage Mapper algorithm on the A375 and CTC datasets using their native time resolution. We evaluate these methods using the AUC, the F1-score, the Precision, the Recall and the Accuracy.

| Method | Dataset | AUC | F1 | Precision | Recall | Accuracy |
| --- | --- | --- | --- | --- | --- | --- |
| **Lineage Mapper** | **A375 (ours)** | - | 0.1178 | 0.1017 | 0.2246 | - |
|  | **Fluo-N2DH-SIM+ (CTC)** | - | 0.7997 | 0.9791 | 0.6899 | - |
|  | **Fluo-N2DL-HeLa (CTC)** | - | 0.1127 | 0.0612 | 0.7666 | - |
|  | **PhC-C2DL-PSC (CTC)** | - | 0.1857 | 0.1096 | 0.8389 | - |
| **Contrastive Division Detection (ours)** | **A375 (ours)** | 0.3641 | 0.4692 | 0.6324 | 0.4018 | 0.9550 |
|  | **Fluo-N2DH-SIM+ (CTC)** | 0.5291 | 0.541 | 0.46740 | 0.655 | 0.9360 |
|  | **Fluo-N2DL-HeLa (CTC)** | 0.7527 | 0.7757 | 0.8648 | 0.7032 | 0.9932 |
|  | **PhC-C2DL-PSC (CTC)** | 0.6106 | 0.7023 | 0.6962 | 0.7084 | 0.9873 |

On the A375 dataset, our method significantly outperformed LM, achieving an F1-score of 0.47 versus Lineage Mapper’s 0.11. However, the comparison yielded mixed results across the CTC datasets, each with distinct characteristics. For the Fluo-N2DH-SIM+ dataset, which contains simulated nuclei of HL60 cells stained with Hoechst,their simulated nature seemingly impact the performance of our method negatively, leading to LM outperforming our approach. The lack of significant evolution in cell appearance and phenotype within this dataset renders our method less effective than LM. Conversely, in the PhC-C2DL-PSC dataset, which comprises pancreatic stem cells imaged with Phase Contrast microscopy on a polystyrene substrate, our method surpassed LM, registering an F1-score of 0.7 against LM’s 0.18. Similarly, on the Fluo-N2DL-HeLa dataset, which closely resembles our own dataset, our method again outperformed LM, achieving an F1-score of 0.77 compared to LM’s 0.11. These notable improvements on the latter two datasets can be attributed to the distinct cell characteristics captured by morphology or fluorescent expression, enabling our model to more effectively discern division events even under challenging conditions.

Additionally, our method demonstrated the highest performance on the HeLa dataset, which is not only the most similar to our own dataset but also has the lowest temporal resolution (30 minutes) among the three evaluated CTC datasets.

We also investigate the performance of our method as a function of the temporal resolution. To this end, we conduct an experimental time subsampling on the CTC datasets in our benchmarking, to observe the variation in our algorithm’s performance with changing temporal resolutions and to align these conditions with those of our dataset. This process involves adjusting the temporal resolution of the dataset to time-step sizes of 60, 120, 180 and 240 minutes. Our analysis of these results focuses however mostly on time intervals up to 180min, as beyond this time resolutions, results are difficult to interpret due to the very low number of remaining frames in the videos after the time subsampling. The results are partially presented in Table 4 and presented in full in Appendix A of the Supplementary Materials. This analysis reveals that our method maintains a coherent and satisfactory level of performance when subjected to significant reductions in temporal resolution. In the case of the HeLa dataset we produce classification scores well above commonly-used unsupervised methods. This evaluation provides insights into the performance of our method, especially at low temporal resolutions on benchmarking datasets. For the Fluo-N2DH-SIM+ dataset, LM outperforms our method, which can be attributed to the limited motion in the simulated dataset and the previously noted static cell appearance. On the Fluo-N2DL-HeLa and PhC-C2DL-PSC datasets, LM’s performance yields F1-scores ranging between 0.08 and 0.18, displaying significantly low recalls across all temporal resolutions. Conversely, our method experiences a performance decrease from the native time resolution results, which correlates with the decreased temporal resolutions. Nonetheless, the performance remains satisfactory, with F1-scores of 0.69 at Δ = 60*min*, 0.64 at Δ = 120*min*, and 0.46 at Δ = 180*min*, alongside precision scores of 0.85, 0.79, and 0.53 respectively. For the PhC-C2DL-PSC dataset, the F1-score was relatively stable across various time resolutions, ranging between 0.25 and 0.30.

**Table 4.** Benchmarking of our proposed division detection method against the Lineage Mapper algorithm on the CTC datasets using artificial time resolutions of 60min, 180min and 240min. We evaluate the two methods at these time-steps using the the F1-score, the Precision, the Recall. This table is excerpted from the table in Appendix A of the Supplementary Materials which contains evaluation of the 120min time resolution, as well as the precision-recall AUC and accuracy metrics, for all time resolutions.

| Dataset | Method | F1-score |  |  | Precision |  |  | Recall |  |  |
| --- | --- | --- | --- | --- | --- | --- | --- | --- | --- | --- |
|  |  | 60min | 180min | 240min | 60min | 180min | 240min | 60min | 180min | 240min |
| Fluo-N2DH-SIM+ (CTC) | LM | 0.8291 | 0.7403 | 0.6410 | 0.9807 | 1.0 | 1.0 | 0.7261 | 0.5881 | 0.4722 |
|  | CDD | 0.3859 | 0.2857 | 0.2435 | 0.3548 | 0.2592 | 0.2176 | 0.42307 | 0.3181 | 0.2765 |
| Fluo-N2DL-HeLa (CTC) | LM | 0.1577 | 0.1834 | 0.2568 | 0.7255 | 0.8156 | 0.7232 | 0.0890 | 0.1045 | 0.1574 |
|  | CDD | 0.6933 | 0.4675 | 0.4420 | 0.8524 | 0.5373 | 0.5145 | 0.5842 | 0.4137 | 0.3875 |
| PhC-C2DL-PSC (CTC) | LM | 0.1211 | 0.0841 | 0.5740 | 0.7740 | 0.6268 | 0.5740 | 0.0679 | 0.0465 | 0.02794 |
|  | CDD | 0.3007 | 0.2499 | 0.3962 | 0.3125 | 0.3611 | 0.3962 | 0.2898 | 0.1911 | 0.1584 |

To sum up, the self-supervised features with custom augmentations as described in Section 3.3.2 as well as the use of the time-context significantly enhance cell division detection across various datasets and outperform traditional methods.

### 4.2. Cell tracking

#### 4.2.1. Tracking features

In this section, we evaluate the efficacy of both individual and combined feature sets within our cell tracking methodology. The performance of these feature sets, as demonstrated in Table 5, is evaluated using the tracking feature metrics introduced insection 3.6.2. We partially present our results in Table 5, with the full results table in Appendix B of the Supplementary Materials integrating every metrics and benchmarking additional features and various weighted combinations of the feature set.

**Table 5.** Evaluation of the performance of individual tracking features and combination of tracking features. We assess each individual features and various feature combination using our dataset (A375 at a time resolution of 240min) using matching cell pairs from our ground truth. With each feature - when used - we indicate in between parenthesis the weight used for this features in the linear combination. We assess the performance of a feature or feature set using the Top-1, Top-3 and Mean Rank metrics both across the whole frame and within a limited distance range. Our complete evaluation is available in the Appendix B of the Supplementary Materials - it includes evaluation through all of our metrics as well as additional features and feature combinations.

|  | Feature Type | Feature | Top-1<br>(whole frame) | Top 1<br>(distance range) | Mean Rank<br>(whole frame) | Mean Rank<br>(distance range) |
| --- | --- | --- | --- | --- | --- | --- |
| Individual<br>Features | Hand-Crafted<br>Features | Overlap | 0.691 | 0.724 | 4.293 | 0.720 |
|  |  | Centroid | **0.776** | **0.776** | **0.403** | **0.381** |
|  |  | Size | 0.117 | 0.436 | 9.456 | 1.495 |
|  | Deep Features<br>(ImageNet) | alexnet | 0.266 | 0.497 | 6.871 | 1.059 |
|  |  | resnet | 0.206 | 0.522 | 8.832 | 1.282 |
|  | Deep Features<br>(ours) | division-simCLR | 0.333 | 0.572 | 5.912 | 1.088 |
|  |  | division-simCLR<sup>3</sup> | 0.569 | 0.672 | 3.817 | 0.7585 |
|  |  | tracking-simCLR | 0.630 | 0.679 | 1.349 | 0.643 |
| Weighted<br>Features<br>Combination | Lineage Mapper<br>Features | Overlap (1.0) +<br>Centroid (0.5) +<br>Size (0.2) | 0.774 | 0.778 | 0.480 | 0.400 |
|  | Deep<br>Features | tracking-simCLR (0.15) +<br>centroid (1.0) | **0.847** | **0.847** | 0.330 | 0.297 |
|  | Deep and Hand-Crafted<br>Features Combination | Centroid (1.0) +<br>Overlap (0.1) +<br>tracking-simCLR (0.15) +<br>division-simCLR<sup>3</sup> (0.15) +<br>resnet<sup>3</sup> (0.05) | **0.847** | **0.847** | **0.313** | **0.277** |

The analysis reveals that the centroid feature is the most effective in our dataset, achieving a top-1 match in 77% of cases. While this performance aligns with expectations, it also underscores the need for additional spatial features to enhance match accuracy in our graph-based tracking algorithm. In contrast, the size feature, a key component of the Lineage Mapper algorithm, demonstrates limited effectiveness, particularly in whole-frame scenarios (top-1 score of 0.11) and improves slightly within the distance range (top-1 score of 0.43). This suggests that rapid morphological changes between frames and a diversity of cell morphologies within a cell’s proximity are challenging the efficacy of the size feature.

Deep features, particularly those incorporating time-context, outperform the size feature in all metrics. Notably, when combined with spatial features (as in the ‘ImageNet’ Features set), these deep features achieve a top-1 match of 78% in whole-frame scenarios and 80% within the distance range, surpassing both the spatial and Lineage Mapper feature sets. In terms of metrics relevant to our approach (with constraints applied to a graph) like top-3 in range, these combined features maintain similar high scores, around the mid 90%. Our custom deep features, including division-simCLR and tracking-simCLR, demonstrated robust performance on their own. The tracking-simCLR feature, in particular, achieves a top-1 score of 63% in the whole frame and 67% within the distance range, with top-3 scores reaching 85% and 92%, respectively. These results position it as the most effective non-spatial feature. When integrated with both spatial and deep features, this combination yielded the highest top-1 score of 84% across both whole frame and distance range scenarios.

Overall, our assessment underscores the notable performance of the features introduced in this work, particularly those developed using the simCLR framework and notably the one using a weakly-supervised approach. We categorize and evaluate feature sets based on their characteristics and the data availability in our evaluation datasets. Two primary sets of features emerge as focal points for an in-depth evaluation:

##### 1. Standa4rd Features

These are the hand-crafted features, which are formulated based on methodologies described in the Lineage Mapper study[7].

##### 2. Deep Features

This set comprises a blend of deep learning-based features and hand-crafted features. It includes simCLR features, contingent upon the availability of adequate model. In scenarios where sufficient data was present to train a model, we utilize a combination of deep and hand-crafted features, as detailed in the ‘Deep and Hand-Crafted Feature Combination’ in Table 5. In cases with more limited training data availability, we employ the ‘ImageNet Feature set’ from the same Table, replacing our trained features with pretrained deep features.

This structured evaluation approach allows us to discern the individual and aggregated strengths of various feature sets, providing a comprehensive understanding of their contribution to the overall effectiveness of our cell tracking methodology and deciding of their inclusion for further evaluation.

#### 4.2.2. Cell tracking method comparison

We evaluate the performance of various configurations of our cell tracking algorithm, each employing distinct sets of features. This comparative analysis spans multiple methodologies:

##### 1. Lineage Mapper

This method serves as our benchmark for unsupervised tracking [7], providing a baseline against which we compare our developed algorithms.

##### 2. Global Tracking

In this variant, we deploy our graph-based tracking algorithm devoid of the rolling window and Markov Decision Process (MDP) elements. Here, tracking is conducted on the entire graph 𝒢 simultaneously, utilizing the shortest mean path algorithm for path determination.

##### 3. Sliding Window Assignment Graph (SWAG)

This configuration of our method experiments with different window sizes, exploring their impact on tracking efficacy.

##### 4. Pyramidal Sliding Window Assignment Graph (P-SWAG)

This approach combines multiple sliding window sizes concurrently. The resultant tracklets are then integrated into the MDP algorithm, aiming to convey the algorithm’s tracking ability by leveraging a multi-scale perspective.

For the evaluation of these methodologies, we apply the tracking performance metrics as defined in section 3.6.2. The results are presented in Table 6.

**Table 6.**
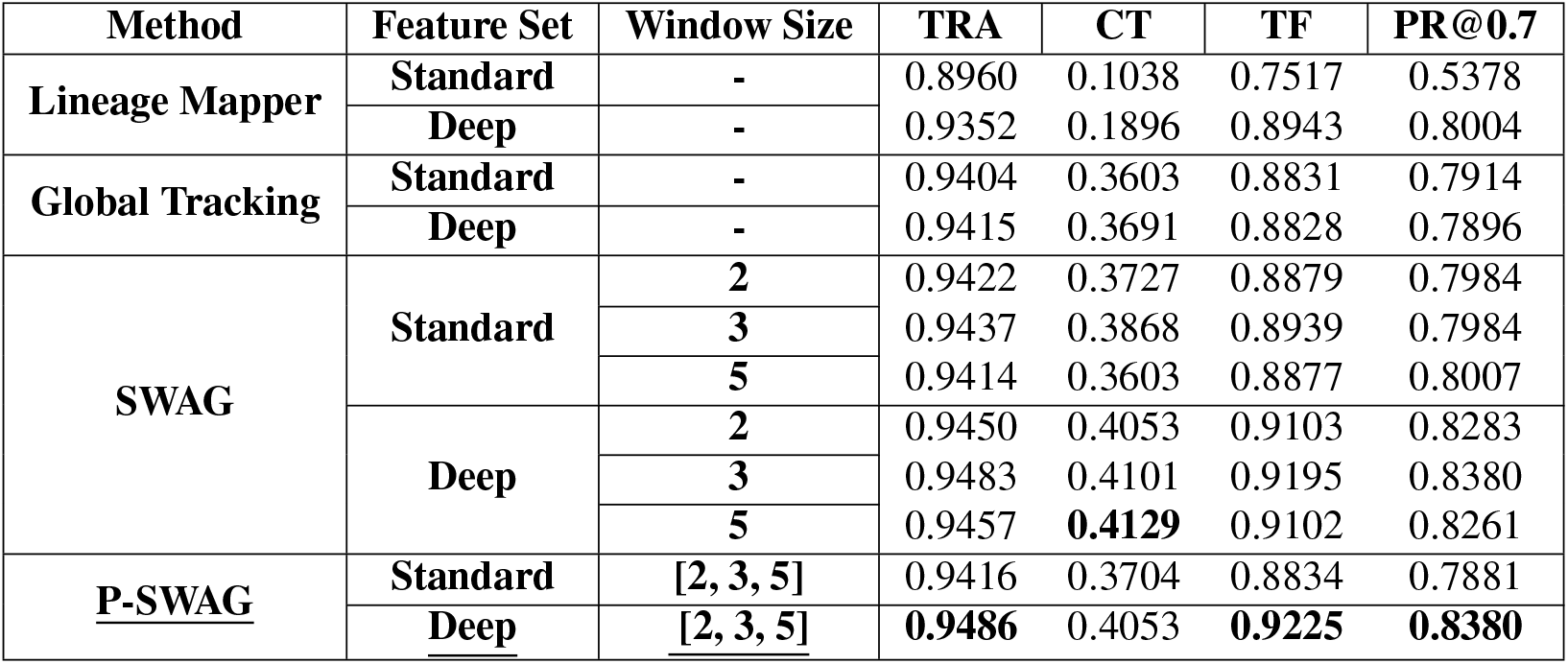
Comparison of the tracking performance of Lineage Mapper with methods introduced in this work on our dataset (A375 at a time resolution of 240min) in combination with different feature sets (Standard and Deep). For the sliding window tracking algorithm (SWAG and P-SWAG) we also showcase different window sizes. For each model configuration we assess the performance using the TRA, the Complete Tracks (CT), the Track Fraction (TF) and the Partial Reconstruction at 70% (PR@0.7).

| Method | Feature Set | Window Size | TRA | CT | TF | PR@0.7 |
| --- | --- | --- | --- | --- | --- | --- |
| Lineage Mapper | Standard | - | 0.8960 | 0.1038 | 0.7517 | 0.5378 |
|  | Deep | - | 0.9352 | 0.1896 | 0.8943 | 0.8004 |
| Global Tracking | Standard | - | 0.9404 | 0.3603 | 0.8831 | 0.7914 |
|  | Deep | - | 0.9415 | 0.3691 | 0.8828 | 0.7896 |
| SWAG | Standard | 2 | 0.9422 | 0.3727 | 0.8879 | 0.7984 |
|  |  | 3 | 0.9437 | 0.3868 | 0.8939 | 0.7984 |
|  |  | 5 | 0.9414 | 0.3603 | 0.8877 | 0.8007 |
|  | Deep | 2 | 0.9450 | 0.4053 | 0.9103 | 0.8283 |
|  |  | 3 | 0.9483 | 0.4101 | 0.9195 | 0.8380 |
|  |  | 5 | 0.9457 | **0.4129** | 0.9102 | 0.8261 |
| P-SWAG | Standard | [2, 3, 5] | 0.9416 | 0.3704 | 0.8834 | 0.7881 |
|  | Deep | [2, 3, 5] | **0.9486** | 0.4053 | **0.9225** | **0.8380** |

Table 6 shows that our methods outperform Lineage Mapper across all configurations on our dataset, with P-SWAG achieving the highest TRA of 0.95. Yet, metrics like Complete Tracks (CT), Track Fraction (TF), and Partially Reconstructed at 70% (PR@0.7) provide more relevant insights for cell biology applications, emphasizing accuracy in track reconstruction. For those metrics, our deep feature set consistently outperforms the standard set, and P-SWAG shows the best results with stable performance across various window sizes.

Incorporating deep features into Lineage Mapper improves its scores, e.g., TF from 0.88 to 0.92 and PR@0.7 from 0.78 to 0.83. However, our method, especially with the sliding window approach, surpassed Lineage Mapper under every configuration, highlighting the robustness of our tracking strategy even with basic features.

Upon these results, P-SWAG with deep features stands out as the most effective configuration to accurately track cellular movements and interactions.

#### 4.2.3. Tracking benchmarking

In this part of our study, we undertake a comparative assessment of our tracking methodology against the Lineage Mapper algorithm [7]. Our results are presented in Table 7 as well as in Appendix C of the Suplemental Materials. Table 7 compares our dataset and the CTC datasets’ tracking using Lineage Mapper and our method, whereas the Table in the Supplementary Materials provides the benchmarking through combination of methods and feature sets.

**Table 7.**
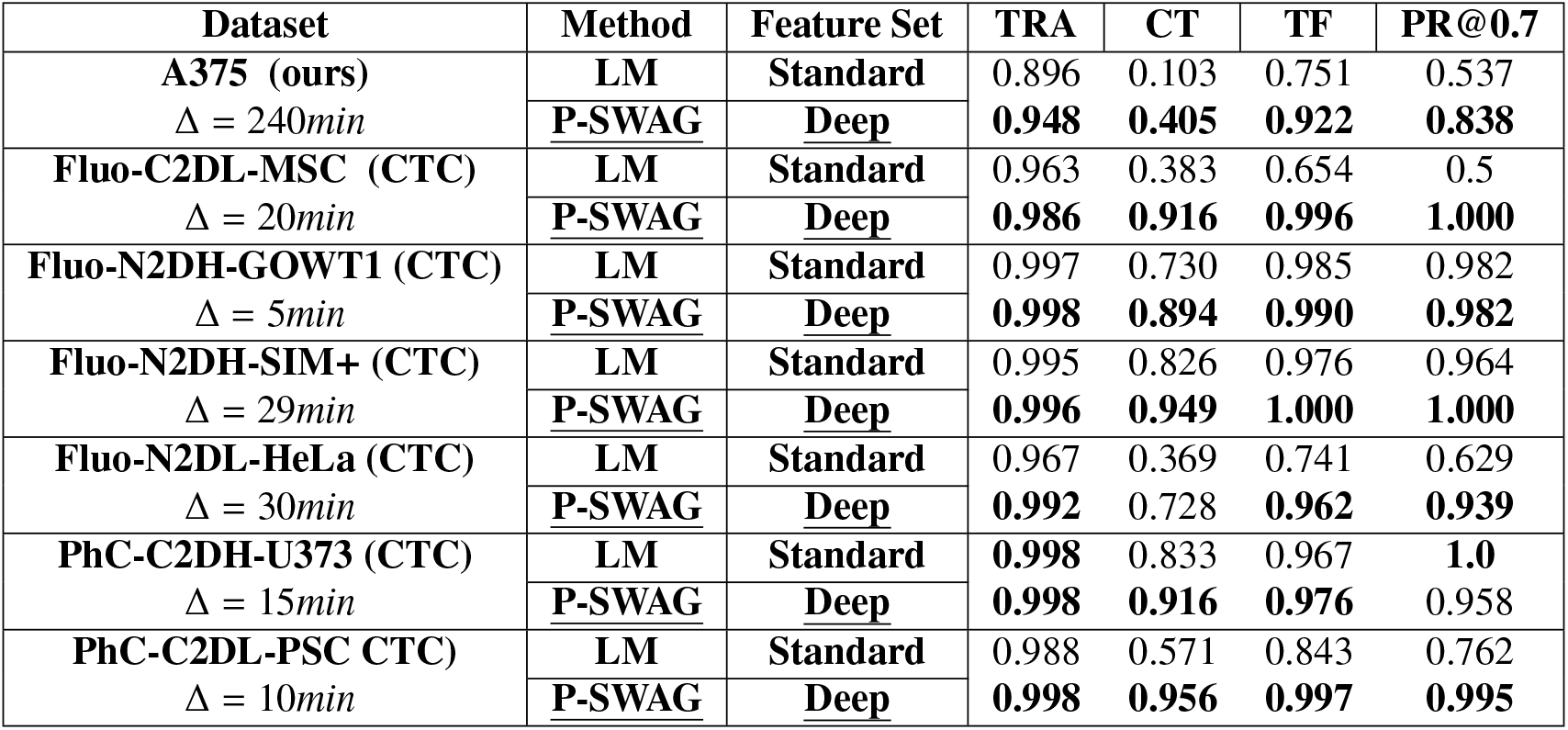
Benchmarking of our tracking method Pyramidal Sliding Window Assignment Graph (P-SWAG) with Lineage Mapper (LM) on two sets of tracking features (Standard Feature) on our dataset (A375) and the CTC datasets in their native time resolution. For each dataset and model we assess the performance using the TRA, the Complete Tracks (CT), the Track Fraction (TF) and the Partial Reconstruction at 70% (PR@0.7).

Our evaluation indicates that our method generally exceeds the performance of the Lineage Mapper across different datasets. Notably, instances where the performance appears equivalent are often those where both methods achieve near-optimal results. Our method consistently outperforms Lineage Mapper, particularly when deep features are used, though there were exceptions. For instance, in the Fluo-N2 DH-GOWT1 dataset, deep features do not improve outcomes, likely due to the dataset’s short 5-minute time steps and minimal frame-to-frame changes. Similarly, in the PhC-C2DH-U373 dataset, neither our approach nor deep features significantly outdid Lineage Mapper, possibly due to its phase contrast imaging and high time-resolution, where deep features might be less effective.

A critical aspect of our evaluation involves addressing the discrepancy in temporal resolution between our dataset and the CTC dataset. To this end, we conduct performance assessments over a range of time-step sizes, from 60 to 240 minutes, using a time-subsampling method. This aspect of the evaluation is pivotal in demonstrating the flexibility and efficiency of our method under varying conditions and temporal resolutions. We thus adjust the temporal resolution of CTC datasets (60 to 180 minutes increments) to assess our tracking robustness to such changes. The results presented in Table 8 and more completely in Appendix D of the Supplementary Materials, indicate that our approach generally performs satisfyingly in scenarios characterized by altered temporal resolutions. Indeed, our method maintains superior performance over Lineage Mapper, particularly in challenging conditions like the PhC-C2DH-U373 dataset, showing our approach’s robustness at various temporal resolutions. Notably, our performance remains high (*TRA* = 0.995, *CT* = 0.81, *TF* = 0.97, *PR*@0.7 = 0.95 at Δ = 180*min*) even when Lineage Mapper’s declines. However, the simulated Fluo-N2DH-SIM+ dataset did not see benefits from deep features, similar to our findings in division detection. These results underline our method’s effectiveness and adaptability across different datasets and conditions showcasing its potential to address the challenges posed by diverse tracking scenarios in low temporal resolution microscopy.

**Table 8.**
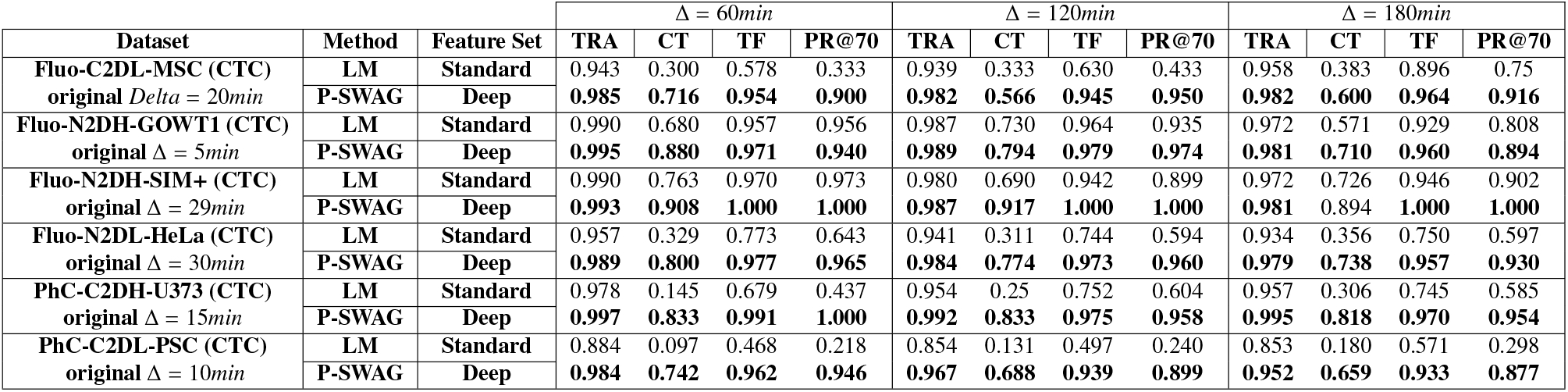
Benchmarking of our tracking method Sliding Window Assignment Graph (SWAG) with Lineage Mapper (LM) on the CTC datasets using artificial temporal resolutions of 60min, 120min and 180min. We evaluate the two methods at these time-steps using the TRA, the Complete Tracks (CT), the Track Fraction (TF) and the Partial Reconstruction at 70% (PR@0.7). Additional results for this evaluation are available in Appendix D of the Supplementary Materials.

For our qualitative analysis, we employ two types of visualizations: dendrograms and track visualizations. The dendrograms presented in Figure 10 highlight the differences in tracking outcomes between our method and the Lineage Mapper method. They illustrate how our method more precisely reconstructs track durations and points of entry and exit, aligning more closely with the actual observed data. Although our approach generally provides more accurate results, it occasionally extends some tracks beyond their actual stopping points. This is likely due to limitations in the sliding window approach used to determine when tracks should end when faced with either low density or high similarity of neighboring cells. In addition to dendrograms, we use tracking examples to further highlight the differences between our method and the Lineage Mapper. The first example in Figure 11 shows how our method continues to track a cell even after the ground truth ceases to do so, thanks to the reliance on strong feature similarities for determining the continuation of tracks. On the other hand, the Lineage Mapper completely loses track of the cell after the third frame. In the second example on Figure 11, our method precisely matches the ground truth by accurately tracking a cell through its final four frames, whereas the Lineage Mapper inaccurately attributes the cell’s origin to a wrong starting point. These examples underscore our method’s capability to robustly identify cell matches despite their morphological and phenotypical changes from frame to frame, adeptly handling both subtle and abrupt changes. Conversely, the Lineage Mapper often incorrectly merges tracks by linking cells based on features that are inconsistently relevant in scenarios of low temporal resolution tracking.

**Figure 10.**
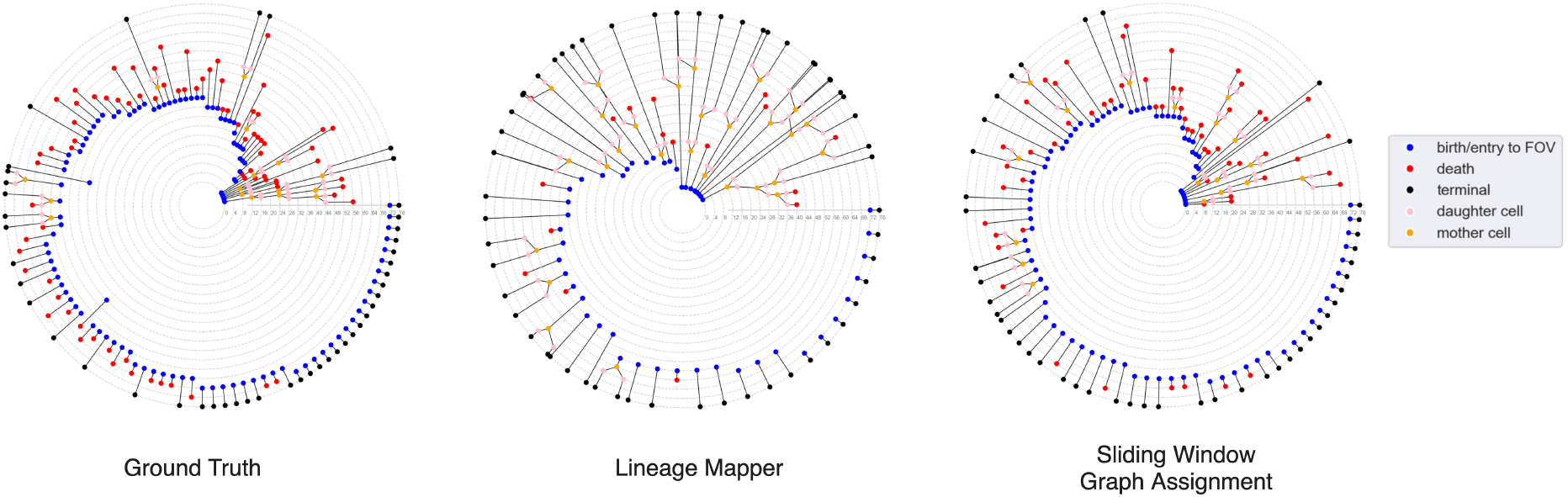
Radial dendrograms for qualitative evaluation of track reconstruction quality. Each ring in the dendrogram corresponds to a frame, starting from the center, and these visualizations distinctly highlight the duration, lineage, and entry/exit points of individual cells. A legend explains the types of nodes shown. We display three versions for the same video: the ground truth, tracks from the Lineage Mapper, and results from our method using the Sliding Window Assignment Graph (SWAG) method. SWAG achieves more accurate track reconstructions compared to the Lineage Mapper, which tends to overmerge tracks resulting in fewer, longer tracks. In contrast, SWAG aligns more closely with the ground truth, although some tracks exhibit slight over-persistence, leading to occasional merges, especially in very short tracks.

**Figure 11.**
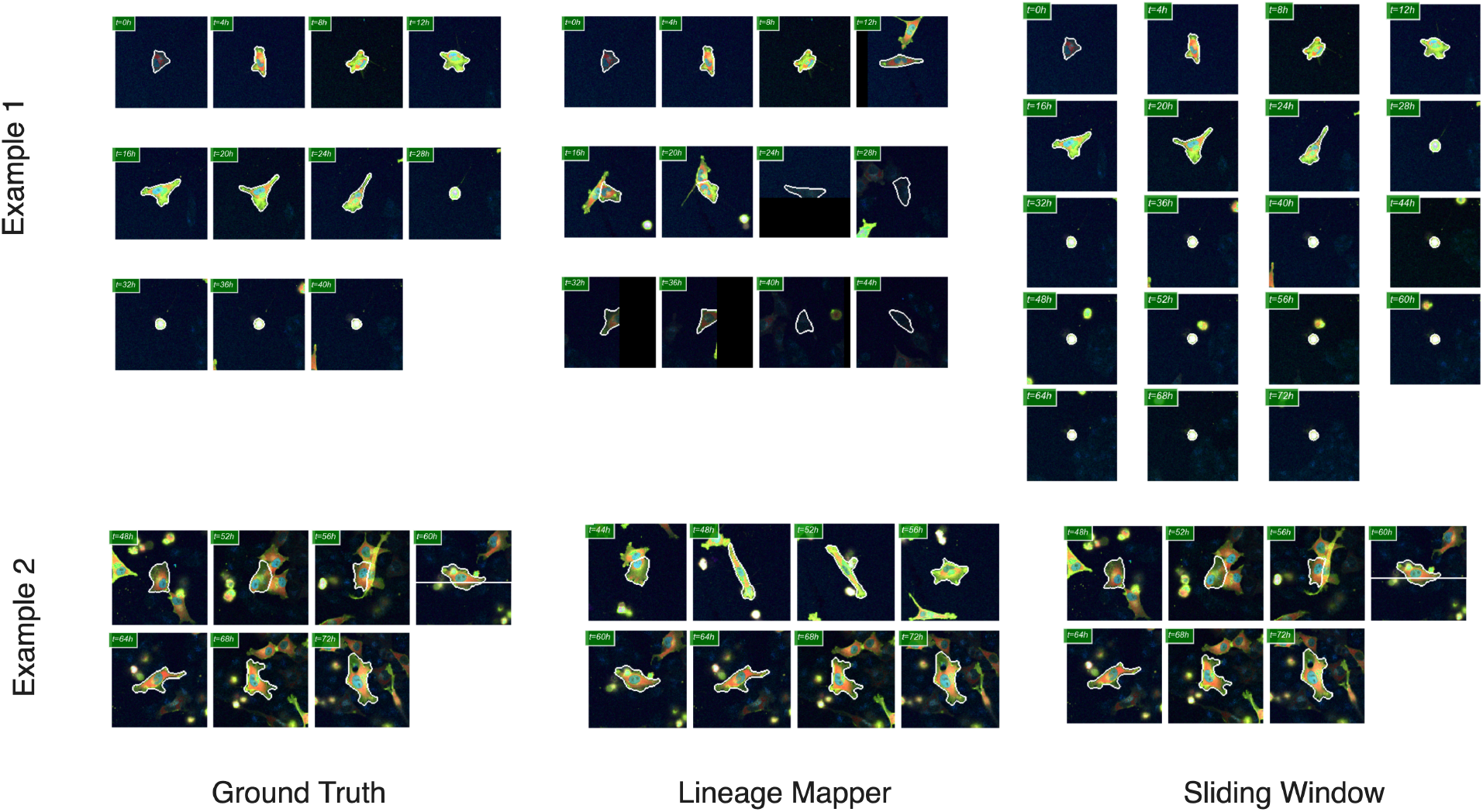
Two examples of track reconstructions from video analysis. The first example illustrates tracks originating from a single cell, highlighting how each method captures the initial movements of the cell. The second example demonstrates tracks converging to end at the same cell, showcasing how each method manages the conclusion of tracks. These scenarios are compared against the ground truth and the results from the Lineage Mapper and our Sliding Window Assignment Graph (SWAG) method. The examples reveal that the Lineage Mapper tends to overmerge tracks by using a features and track assignment method not well-suited for low temporal resolution tracking. In contrast, SWAG effectively handles both subtle and abrupt changes from frame to frame, maintaining robust tracking accuracy in these challenging conditions.

## 5. Discussion

Time-lapse data plays a crucial role in drug screening, particularly for observing the effects of treatments over extended periods of time. In particular, live-cell High Content Screening (HCS) assays produce video-microscopy data that connects drug-induced single-cell phenotypes to morphological changes and molecular signals over time, using fluorescent markers to highlight relevant signaling pathways. To derive richer and more meaningful insights and biological interpretations from live-cell HCS data, accurately tracking individual cells over time is essential. Indeed, this tracking enables the study of cellular morphological and phenotypic changes under treatment influence. However, long-term imaging presents challenges, such as phototoxicity, resulting from frequent exposure to light. To mitigate this, temporal resolution must be reduced, complicating the tracking of individual cells and their lineage across time. Our paper introduces methods enabling to accurately detect cell division events and track cells over extended periods of time, even with lower temporal resolution. This capability increases the range of possible experiments and thus the biological insights one can generate in the subsequent downstream analysis, allowing for precise profiling of cellular responses to treatments over extended periods of time.

Indeed, studying live individual cells over extended periods might significantly benefit to our understanding of cellular behavior and drug interactions. This approach not only allows us to dynamically monitor how cells grow, divide, and interact but also offers a detailed look at how they respond to various treatments. By observing these processes in real-time, we gain valuable insights into the natural progression of cellular activities and the nuanced mechanisms of drug effects, including their efficacy and potential toxicity. Long-term live-cell imaging helps us to track the activation of signaling pathways in response to stimuli, revealing how cells communicate and react at the molecular level. It also sheds light on the diversity within cell populations, uncovering different phenotypic responses to treatments. This is crucial for understanding why drugs might work effectively in some cells but not in others. Furthermore, such detailed observation helps us explore the intricacies of the cell cycle, apoptosis, and intracellular trafficking - processes that are inherently dynamic. Thanks to the research we have presented and by extending it to downstream analysis, we might see firsthand how cells progress through various growth phases, how they program themselves for death, and how they transport components internally. Such knowledge might be key to developing new anticancer therapies or more generally understanding cellular organization. Moreover, observing how cells interact with each other and their surroundings gives us clues about tissue organization and disease progression. By integrating these observations into predictive models, we might better anticipate cell responses to different treatments, paving the way for more effective and safer therapeutic strategies. Our paper thus proposes a way to push the frontiers of live-cell assay designs by enabling the study of video-microscopy data with low temporal resolution.

Furthermore, our research also highlights the role of contrastive and Self-Supervised Learning (SSL) in generating features useful for the tasks of division detection and tracking. As demonstrated in this work, SSL features enable us to harness extensive fluorescent video-microscopy datasets with minimal or no human annotations. We have demonstrated the effectiveness of applying specific augmentations tailored to particular tasks such as cell division detection and tracking. Notably, in tracking, we have enhanced the SSL model by incorporating time-based augmentations, teaching the model to recognize cell changes over different time points, based on weakly supervised annotations. When comparing these time-based augmentations to other SSL features presented in our study, we observed significant improvements in performance on target tasks. This underscores the value of including task-specific augmentations and, more broadly, the benefits of utilizing time-based SSL representations for cellular analysis. Indeed, the methods and features we propose are suitable beyond cell division detection and tracking for the analysis of individual cells within drug-screening studies, particularly where annotations are scarce and expensive with varying experimental conditions. Within such studies, using time-based SSL features might reinforce our capacity to predict and comprehend how cells respond to drug interventions over time and more generally our ability to extract rich, dynamic insights from volumes of un-annotated data, in turn enhancing our predictions regarding drug interactions and cellular targets.

## 6. Conclusion

This study presents a comprehensive workflow for cell division detection and tracking in live cell microscopy, addressing challenges posed by low temporal resolution and limited expert annotations. By integrating advanced contrastive learning techniques with graph-based approaches, we have developed methodologies that significantly enhance the accuracy and robustness of dynamic cellular process analysis.

For cell division detection, the employment of a simCLR model trained on sequential image patches enables precise identification of morphological changes indicative of cell division. This approach, substantiated by improved F1-scores, underscores the effectiveness of incorporating temporal context into the division detection process.

For cell tracking, the development of features derived from weakly supervised learning and temporal augmentations facilitates the robust representation of cellular dynamics. Coupled with our Sliding Window Assignment Graph (SWAG) technique, they demonstrate state-of-the-art performances in key tracking metrics, including Track Reconstruction Accuracy, Complete Tracks, and Track Fraction. The methodology’s adaptability to varying temporal resolutions is particularly advantageous for extended live-cell drug screening studies using video-microscopy, offering a means to circumvent the challenges of photo-toxicity that leads to a decrease of temporal resolution.

Our findings, benchmarked against unsupervised methods and established datasets, reveal a consistent improvement in detecting and tracking cellular entities, even in challenging imaging conditions. The adaptability of our approach to different temporal resolutions and its effectiveness in mitigating photo-toxic effects highlight its potential. This work contributes to the growing body of knowledge in cellular imaging and analysis, and opens avenues for more accurate and efficient studies in cell biology, particularly in contexts where temporal resolution and photo-toxicity are critical factors.

## Acknowledgments

This work was supported by grants from Région Ile-de-France through the DIM ELICIT program. The data analysis was carried out using HPC resources from GENCI-IDRIS (Grant 2023—AD011011156R3).

We sincerely thank Catherine Lacayo and Laura Savy for contributing to the development the cell biology tools and conducting the assays that provide the data used in this work, as well as Payman Amiri for his constructive feedback and thoughtful comments regarding this work.

