## Supplementary Materials for "Contrastive learning for cell division detection and tracking in live cell imaging data"

Available online at www.sciencedirect.com

ScienceDirect

Pattern Recognition 00 (2024) 1–4

Pattern Recognition

### Contrastive learning for cell division detection and tracking in live cell imaging data - Supplementary Materials

Daniel Zyss[a,b,c,d], Amritansh Sharma[d], Susana A Ribeiro[d], Claire E Repellin[e], Oliver Lai[f], Mary J C Ludlam[d], Thomas Walter[a,b,c,*], Amin Fehri[d]

*[a]Center for Computational Biology (CBIO), Mines Paris, PSL University, Paris, France*
*[b]Institut Curie, PSL University, 75248 Paris Cedex, France*
*[c]INSERM, U900, 75248 Paris Cedex, France*
*[d]Cairn Biosciences Inc, San Francisco, CA 94107, USA*
*[e]ORIC Pharmaceuticals, South San Francisco, CA 94080, USA*
*[f]Revolution Medicines, Redwood City CA 94063, USA*

#### Appendix A. Time-Adjusted Division Detection Benchmarking

| | | Delta=60min | | | | | Delta=120min | | | | | Delta=180min | | | | | Delta=240min | | | | |
|---|---|---|---|---|---|---|---|---|---|---|---|---|---|---|---|---|---|---|---|---|---|
| **Dataset** | **Method** | **AUC** | **F1** | **P** | **R** | **A** | **AUC** | **F1** | **P** | **R** | **A** | **AUC** | **F1** | **P** | **R** | **A** | **AUC** | **F1** | **P** | **R** | **A** |
| **Fluo-N2DH-SIM+ (CTC)** | **LM** | - | 0.8291 | 0.9807 | 0.7261 | - | - | 0.7913 | 0.9827 | 0.7133 | - | - | 0.7403 | 1.0 | 0.5881 | - | - | 0.6410 | 1.0 | 0.4722 | - |
| | **CDD** | 0.3171 | 0.3859 | 0.3548 | 0.42307 | 0.825 | 0.3498 | 0.4358 | 0.3207 | 0.68 | 0.8196 | 0.2440 | 0.2857 | 0.2592 | 0.3181 | 0.8309 | 0.1875 | 0.2435 | 0.2176 | 0.2765 | 0.7845 |
| **Fluo-N2DL-HeLa (CTC)** | **LM** | - | 0.1577 | 0.7255 | 0.0890 | - | - | 0.1756 | 0.8564 | 0.0999 | - | - | 0.1834 | 0.8156 | 0.1045 | - | - | 0.2568 | 0.7232 | 0.1574 | - |
| | **CDD** | 0.7132 | 0.6933 | 0.8524 | 0.5842 | 0.9866 | 0.5339 | 0.6482 | 0.7966 | 0.5465 | 0.9757 | 0.2973 | 0.4675 | 0.5373 | 0.4137 | 0.9785 | 0.2456 | 0.4420 | 0.5145 | 0.3875 | 0.9791 |
| **PhC-C2DL-PSC (CTC)** | **LM** | - | 0.1211 | 0.7740 | 0.0679 | - | - | 0.1130 | 0.75 | 0.0627 | - | - | 0.0841 | 0.6268 | 0.0465 | - | - | 0.0532 | 0.5740 | 0.02794 | - |
| | **CDD** | 0.1736 | 0.3007 | 0.3125 | 0.2898 | 0.9699 | 0.1248 | 0.2803 | 0.375 | 0.2238 | 0.9758 | 0.1378 | 0.2499 | 0.3611 | 0.1911 | 0.9682 | 0.1274 | 0.2263 | 0.3962 | 0.1584 | 0.9501 |

Table A.1. Benchmarking of our proposed division detection method against the Lineage Mapper algorithm on the CTC datasets using artificial time resolutions of 60min, 120min, 180min and 240min. We evaluate the two methods at these time-steps using the the F1-score, the Precision, the Recall, the precision-recall AUC and the Accuracy.

*Corresponding author

#### Appendix B. Tracking Feature Benchmarking

| | Feature Type | Feature | Top-1 (whole frame) | Top-3 (whole frame) | Mean Rank (whole frame) | Top-1 (distance range) | Top-3 (distance range) | Mean Rank (distance range) |
|---|---|---|---|---|---|---|---|---|
| **Individual Features** | **Hand-Crafted Features** | **Overlap** | 0.691 | 0.786 | 4.293 | 0.724 | 0.894 | 0.720 |
| | | **Centroid** | **0.776** | **0.959** | **0.403** | **0.776** | **0.958** | **0.381** |
| | | **Size** | 0.117 | 0.271 | 9.456 | 0.436 | 0.771 | 1.495 |
| | **Deep Features (Image-Net)** | **alexnet** | 0.275 | 0.497 | 7.15 | 0.556 | 0.852 | 1.044 |
| | | **resnet** | 0.231 | 0.394 | 8.522 | 0.512 | 0.798 | 1.292 |
| | | **alexnet (RGB)** | 0.266 | 0.408 | 6.871 | 0.497 | 0.857 | 1.059 |
| | | **resnet (RGB)** | 0.206 | 0.418 | 8.832 | 0.522 | 0.832 | 1.282 |
| | | **alexnet$^3$** | 0.477 | 0.620 | 5.000 | 0.669 | 0.862 | 0.881 |
| | | **resnet$^3$** | 0.408 | 0.556 | 6.822 | 0.615 | 0.847 | 1.00 |
| | **Deep Features (Ours)** | **division-simCLR** | 0.333 | 0.517 | 5.912 | 0.572 | 0.843 | 1.088 |
| | | **division-simCLR$^3$** | 0.569 | 0.706 | 3.817 | 0.672 | 0.885 | 0.785 |
| | | **tracking-simCLR** | 0.630 | 0.857 | 1.349 | 0.679 | 0.921 | 0.643 |
| **Weighted Features Combination** | **Spatial Features** | **Centroid (1.0) + Overlap (1.0)** | 0.778 | 0.958 | 0.390 | 0.778 | 0.957 | 0.368 |
| | **Lineage Mapper Features** | **Overlap (1.0) + Centroid (0.5) + Size (0.2)** | 0.774 | 0.938 | 0.480 | 0.778 | 0.950 | 0.400 |
| | **ImageNet Features** | **alexnet (0.2) + resnet (0.2) + alexnet$^3$ (1.0) + resnet$^3$ (1.0)** | 0.433 | 0.600 | 5.364 | 0.650 | 0.886 | 0.757 |
| | | **alexnet (0.1) + resnet (0.1) + alexnet$^3$ (0.5) + resnet$^3$ (0.5) + Centroid (1.0) + Overlap (0.3)** | 0.783 | 0.881 | 1.231 | 0.802 | 0.931 | 0.430 |
| | **Our Features** | **division-simCLR (0.2) + division-simCLR$^3$ (0.5) + tracking-simCLR (1.0)** | 0.709 | 0.891 | 1.039 | 0.763 | 0.940 | 0.435 |
| | | **division-simCLR (0.05) + division-simCLR$^3$ (0.05) + tracking-simCLR (0.15) + centroid (1.0) + overlap (0.1)** | **0.847** | 0.955 | 0.315 | **0.847** | 0.955 | **0.277** |
| | | **tracking-simCLR (0.15) + centroid (1.0)** | **0.847** | 0.955 | 0.330 | **0.847** | 0.955 | 0.297 |
| | **Deep and Hand-Crafted Feature Combination** | **Centroid (1.0) + Overlap (0.1) + tracking-simCLR (0.15) + division-simCLR$^3$ (0.15) + resnet$^3$ (0.05)** | **0.847** | **0.960** | **0.313** | **0.847** | **0.9605** | **0.277** |

Table B.2. Evaluation of the performance of individual tracking features and combination of tracking features. We assess each individual features and various feature combination using our dataset (A375 at a time resolution of 180min) using matching cell pairs from our ground truth. With each feature - when used - we indicate in between parenthesis the weight used for this features in the linear combination. We assess the performance of a feature or feature set using the Top-1, Top-3 and Mean Rank metrics both across the whole frame and within a limited distance range.

#### Appendix C. Tracking Methods Benchmarking

| Dataset | Method | Feature Set | TRA | CT | TF | PR@0.7 |
|---|---|---|---|---|---|---|
| **A375 (ours)** $\Delta = 180min$ | **LM** | **Standard** | 0.896 | 0.103 | 0.751 | 0.537 |
| | | **Deep** | 0.935 | 0.189 | 0.894 | 0.800 |
| | **P-SWAG** | **Standard** | 0.941 | 0.370 | 0.883 | 0.788 |
| | | **Deep** | 0.948 | 0.405 | 0.922 | 0.838 |
| **Fluo-C2DL-MSC (CTC)** $\Delta = 20min$ | **LM** | **Standard** | 0.963 | 0.383 | 0.654 | 0.5 |
| | | **Deep** | 0.985 | 0.649 | 0.973 | 0.966 |
| | **P-SWAG** | **Standard** | **0.986** | **0.916** | **0.996** | **1.000** |
| | | **Deep** | **0.986** | **0.916** | **0.996** | **1.000** |
| **Fluo-N2DH-GOWT1 (CTC)** $\Delta = 5min$ | **LM** | **Standard** | 0.997 | 0.730 | 0.985 | 0.982 |
| | | **Deep** | **0.998** | 0.817 | 0.989 | 0.991 |
| | **P-SWAG** | **Standard** | **0.998** | **0.894** | **0.990** | **0.982** |
| | | **Deep** | **0.998** | **0.894** | **0.990** | **0.982** |
| **Fluo-N2DH-SIM+ (CTC)** $\Delta = 29min$ | **LM** | **Standard** | 0.995 | 0.826 | 0.976 | 0.964 |
| | | **Deep** | 0.996 | 0.933 | 0.994 | 0.997 |
| | **P-SWAG** | **Standard** | **0.996** | **0.949** | **1.000** | **1.000** |
| | | **Deep** | **0.996** | **0.949** | **1.000** | **1.000** |
| **Fluo-N2DL-HeLa (CTC)** $\Delta = 30min$ | **LM** | **Standard** | 0.967 | 0.369 | 0.741 | 0.629 |
| | | **Deep** | 0.987 | 0.704 | 0.921 | 0.885 |
| | **P-SWAG** | **Standard** | 0.988 | **0.738** | 0.924 | 0.885 |
| | | **Deep** | **0.992** | 0.728 | **0.962** | **0.939** |
| **PhC-C2DH-U373 (CTC)** $\Delta = 15min$ | **LM** | **Standard** | **0.998** | 0.833 | 0.967 | 1.0 |
| | | **Deep** | **0.998** | **0.916** | 0.958 | **0.976** |
| | **P-SWAG** | **Standard** | **0.998** | **0.916** | **0.976** | 0.958 |
| | | **Deep** | **0.998** | **0.916** | **0.976** | 0.958 |
| **PhC-C2DL-PSC CTC)** $\Delta = 10min$ | **LM** | **Standard** | 0.988 | 0.571 | 0.843 | 0.762 |
| | | **Deep** | 0.988 | 0.730 | 0.884 | 0.855 |
| | **P-SWAG** | **Standard** | 0.989 | 0.778 | 0.897 | 0.863 |
| | | **Deep** | **0.998** | **0.956** | **0.997** | **0.995** |

Table C.3. Benchmarking of our tracking method Sliding Window Assignment Grapĥ (SWAG) with Lineage Mapper (LM) on two sets of tracking features (Standard Feature) on our dataset (A375) and the CTC datasets in their native time resolution. For each dataset and model we assess the performance using the TRA, the Complete Tracks (CT), the Track Fraction (TF) and the Partial Reconstruction at 70% (PR@0.7).

#### Appendix D. Benchmarking of Tracking Methods with Adjusted Time Resolutions

| | | | $Δ = 60*min*$ | | | | Δ = 120*min* | | | | Δ = 180*min* | | | | Δ = 240*min* | | | |
|---|---|---|---|---|---|---|---|---|---|---|---|---|---|---|---|---|---|---|
| **Dataset** | **Method** | **Feature Set** | **TRA** | **CT** | **TF** | **PR@70** | **TRA** | **CT** | **TF** | **PR@70** | **TRA** | **CT** | **TF** | **PR@70** | **TRA** | **CT** | **TF** | **PR@70** |
| **Fluo-C2DL-MSC (CTC) original** Δ = 20*min* | **LM** | **Standard** | 0.943 | 0.300 | 0.578 | 0.333 | 0.939 | 0.333 | 0.630 | 0.433 | 0.958 | 0.383 | 0.896 | 0.75 | - | - | - | - |
| | | **Deep** | 0.983 | 0.516 | 0.921 | 0.866 | 0.970 | 0.383 | 0.847 | 0.700 | 0.935 | 0.433 | 0.683 | 0.483 | - | - | - | - |
| | **P-SWAG** | **Standard** | 0.982 | 0.614 | 0.937 | 0.871 | 0.979 | 0.480 | 0.921 | 0.842 | 0.951 | 0.537 | 0.819 | 0.847 | - | - | - | - |
| | | **Deep** | **0.985** | **0.716** | **0.954** | **0.900** | **0.982** | **0.566** | **0.945** | **0.950** | **0.982** | **0.600** | **0.964** | **0.916** | - | - | - | - |
| **Fluo-N2DH-GOWT1 (CTC) original** Δ = 5*min* | **LM** | **Standard** | 0.990 | 0.680 | 0.957 | 0.956 | 0.987 | 0.730 | 0.964 | 0.935 | 0.972 | 0.571 | 0.929 | 0.808 | - | - | - | - |
| | | **Deep** | 0.992 | 0.723 | 0.976 | 0.964 | 0.985 | 0.743 | 0.922 | 0.884 | 0.977 | 0.709 | 0.905 | 0.762 | - | - | - | - |
| | **P-SWAG** | **Standard** | **0.995** | **0.880** | **0.971** | **0.940** | **0.989** | **0.794** | **0.979** | **0.974** | **0.981** | **0.710** | **0.960** | **0.894** | - | - | - | - |
| | | **Deep** | **0.995** | **0.880** | **0.971** | **0.940** | **0.989** | **0.794** | **0.979** | **0.974** | **0.981** | **0.710** | **0.960** | **0.894** | - | - | - | - |
| **Fluo-N2DH-SIM+ (CTC) original** Δ = 29*min* | **LM** | **Standard** | 0.990 | 0.763 | 0.970 | 0.973 | 0.980 | 0.690 | 0.942 | 0.899 | 0.972 | 0.726 | 0.946 | 0.902 | 0.984 | 0.682 | 0.918 | 0.854 |
| | | **Deep** | 0.993 | 0.887 | 0.997 | 0.994 | 0.987 | 0.890 | 0.999 | **1.000** | 0.980 | 0.855 | 0.996 | 0.994 | 0.997 | 0.861 | 0.991 | 0.978 |
| | **P-SWAG** | **Standard** | **0.993** | **0.908** | **1.000** | **1.000** | **0.987** | 0.896 | 0.997 | **1.000** | **0.981** | **0.905** | **1.000** | **1.000** | **0.998** | **0.904** | **0.993** | **0.984** |
| | | **Deep** | **0.993** | **0.908** | **1.000** | **1.000** | **0.987** | **0.917** | **1.000** | **1.000** | **0.981** | 0.894 | **1.000** | **1.000** | **0.998** | **0.904** | **0.993** | **0.984** |
| **Fluo-N2DL-HeLa (CTC) original** Δ = 30*min* | **LM** | **Standard** | 0.957 | 0.329 | 0.773 | 0.643 | 0.941 | 0.311 | 0.744 | 0.594 | 0.934 | 0.356 | 0.750 | 0.597 | 0.973 | 0.497 | 0.878 | 0.811 |
| | | **Deep** | 0.978 | 0.586 | 0.882 | 0.819 | 0.969 | 0.568 | 0.864 | 0.790 | 0.963 | 0.580 | 0.859 | 0.787 | 0.990 | 0.722 | 0.957 | **0.934** |
| | **P-SWAG** | **Standard** | 0.980 | 0.616 | 0.887 | 0.816 | 0.969 | 0.594 | 0.872 | 0.802 | 0.963 | 0.584 | 0.854 | 0.774 | 0.990 | 0.774 | 0.953 | 0.923 |
| | | **Deep** | **0.989** | **0.800** | **0.977** | **0.965** | **0.984** | **0.774** | **0.973** | **0.960** | **0.979** | **0.738** | **0.957** | **0.930** | **0.991** | **0.768** | **0.955** | 0.926 |
| **PhC-C2DH-U373 (CTC) original** Δ = 15*min* | **LM** | **Standard** | 0.978 | 0.145 | 0.679 | 0.437 | 0.954 | 0.25 | 0.752 | 0.604 | 0.957 | 0.306 | 0.745 | 0.585 | 0.956 | 0.232 | 0.775 | 0.556 |
| | | **Deep** | **0.997** | **0.833** | **0.991** | **1.000** | 0.991 | 0.708 | 0.920 | 0.875 | 0.976 | 0.568 | 0.816 | 0.704 | 0.966 | 0.386 | 0.727 | 0.613 |
| | **P-SWAG** | **Standard** | 0.996 | **0.833** | 0.984 | 0.958 | **0.992** | **0.833** | **0.975** | **0.958** | **0.984** | 0.613 | 0.824 | 0.886 | 0.966 | 0.386 | 0.727 | 0.613 |
| | | **Deep** | **0.997** | **0.833** | **0.991** | **1.000** | **0.992** | **0.833** | **0.975** | **0.958** | **0.995** | **0.818** | **0.970** | **0.954** | **0.999** | **0.909** | **1.0** | **1.0** |
| **PhC-C2DL-PSC (CTC) original** Δ = 10*min* | **LM** | **Standard** | 0.884 | 0.097 | 0.468 | 0.218 | 0.854 | 0.131 | 0.497 | 0.240 | 0.853 | 0.180 | 0.571 | 0.298 | 0.936 | 0.374 | 0.773 | 0.562 |
| | | **Deep** | 0.923 | 0.173 | 0.508 | 0.321 | 0.899 | 0.212 | 0.505 | 0.291 | 0.895 | 0.272 | 0.581 | 0.348 | 0.975 | 0.650 | 0.910 | 0.812 |
| | **P-SWAG** | **Standard** | 0.932 | 0.211 | 0.547 | 0.387 | 0.905 | 0.227 | 0.526 | 0.319 | 0.899 | 0.280 | 0.593 | 0.363 | 0.976 | 0.657 | 0.911 | 0.817 |
| | | **Deep** | **0.984** | **0.742** | **0.962** | **0.946** | **0.967** | **0.688** | **0.939** | **0.899** | **0.952** | **0.659** | **0.933** | **0.877** | **0.978** | **0.663** | **0.928** | **0.849** |

Table D.4. Benchmarking of our proposed tracking method Sliding Window Assignment Grapĥ (SWAG) against the Lineage Mapper (LM) algorithm on the CTC datasets using artificial time resolutions of 60min, 120min, 180min and 240min. Some results for the 240min time resolution are omitted due to the resulting duration of the videos being under 5 frames. We evaluate the two methods at these time-steps using the TRA, the Complete Tracks (CT), the Track Fraction (TF) and the Partial Reconstruction at 70% (PR@0.7).
